# Natural Killer cell activation, reduced ACE2, TMPRSS2, cytokines G-CSF, M-CSF and SARS-CoV-2-S pseudovirus infectivity by MEK inhibitor treatment of human cells

**DOI:** 10.1101/2020.08.02.230839

**Authors:** Lanlan Zhou, Kelsey Huntington, Shengliang Zhang, Lindsey Carlsen, Eui-Young So, Cassandra Parker, Ilyas Sahin, Howard Safran, Suchitra Kamle, Chang-Min Lee, Chun Geun Lee, Jack A. Elias, Kerry S. Campbell, Mandar T. Naik, Walter J. Atwood, Emile Youssef, Jonathan A. Pachter, Arunasalam Navaraj, Attila A. Seyhan, Olin Liang, Wafik S. El-Deiry

## Abstract

COVID-19 affects vulnerable populations including elderly individuals and patients with cancer. Natural Killer (NK) cells and innate-immune TRAIL suppress transformed and virally-infected cells. ACE2, and TMPRSS2 protease promote SARS-CoV-2 infectivity, while inflammatory cytokines IL-6, or G-CSF worsen COVID-19 severity. We show MEK inhibitors (MEKi) VS-6766, trametinib and selumetinib reduce ACE2 expression in human cells. In some human cells, remdesivir increases ACE2-promoter luciferase-reporter expression, ACE2 mRNA and protein, and ACE2 expression is attenuated by MEKi. In serum-deprived and stimulated cells treated with remdesivir and MEKi we observed correlations between pRB, pERK, and ACE2 expression further supporting role of proliferative state and MAPK pathway in ACE2 regulation. We show elevated cytokines in COVID-19-(+) patient plasma (N=9) versus control (N=11). TMPRSS2, inflammatory cytokines G-CSF, M-CSF, IL-1α, IL-6 and MCP-1 are suppressed by MEKi alone or with remdesivir. We observed MEKi stimulation of NK-cell killing of target-cells, without suppressing TRAIL-mediated cytotoxicity. Pseudotyped SARS-CoV-2 virus with a lentiviral core and SARS-CoV-2 D614 or G614 SPIKE (S) protein on its envelope infected human bronchial epithelial cells, small airway epithelial cells, or lung cancer cells and MEKi suppressed infectivity of the pseudovirus. We show a drug class-effect with MEKi to stimulate NK cells, inhibit inflammatory cytokines and block host-factors for SARS-CoV-2 infection leading also to suppression of SARS-CoV-2-S pseudovirus infection of human cells. MEKi may attenuate SARS-CoV-2 infection to allow immune responses and antiviral agents to control disease progression.

## Introduction

Coronavirus 2 (SARS-CoV-2) infection progresses to a rapidly lethal adult respiratory distress syndrome (ARDS) associated with high mortality especially among the elderly or those with multiple comorbid conditions [1–5]. Patients with cancer are particularly vulnerable in part due to their weakened immune system and are further at risk due to the immune suppressive effects of chemotherapy [6–8]. The lethality of SARS-CoV-2, the causative agent for the COVID-19 disease, involves a fulminant cytokine storm with bilateral lung infiltrates observed on chest X-rays and CT scans [9]. It has become clear that COVID-19 disease involves multiple organ systems including pulmonary, neurological, renal, hematological and gastrointestinal systems, among others [10–15]. The SARS-CoV-2 virus binds to angiotensin converting enzyme 2 (ACE2) receptors and cellular entry is facilitated by TMPRSS2 protease [16]. Current therapeutic approaches include a number of agents such as anti-inflammatory agents that block IL-6, steroids, anti-viral agents, convalescent serum and alpha receptor blockers [17–21]. There are ongoing approaches for drug discovery and drug repurposing [22, 23].

Once SARS-Cov-2 enters into cells it triggers a host immune response that leads to pathogenesis and disease progression [24]. A SARS-CoV-2 SPIKE protein variant (D614G) has emerged as the dominant pandemic form with evidence that it increases infectivity of the COVID-19 virus [25]. The host inflammatory response phase of COVID-19 is the phase where patients become critically ill leading to high patient mortality [26]. We sought to better understand and modulate the host immune response to SARS-CoV-2 in order to prevent or reduce disease severity. This includes strategies to inhibit expression of ACE2, the receptor SARS-CoV-2 uses to enter cells.

It is clear that while the host systemic inflammatory response makes patients critically ill, the host innate immune system including natural killer (NK) cells is involved in fighting and eliminating virally-infected cells [27]. Over the last 25 years we have studied this innate immune system pathway that the immune system uses to eliminate transformed and cancer cells as well as virally-infected cells [28–34]. Natural killer cells secrete TRAIL which is involved in killing virally-infected as well as transformed cells [35–38]. Thus, our goal was to better understand and modulate the host immune response to increase the innate immune system early in SARS-CoV-2 infection while reducing the severe inflammation that occurs late in the disease course. We further wanted to understand the impact of current therapeutics used to treat COVID-19 on SARS-CoV-2 infectivity factors, the innate immune system and the cellular inflammatory response.

Prior work has suggested that coronavirus SPIKE protein can through ACE2 activate the MAPK pathway and downstream inflammatory responses [39]. Other data suggested that MAPK regulates ACE2 [40], and so we investigated the impact of MEK inhibition on ACE2 expression as a strategy to attenuate early SARS-Cov-2 infection. Since remdesivir has been shown to reduce hospitalization [19] and may reduce mortality in patients with severe COVID-19 infection [41], we hypothesized that suppression of viral entry into cells through inhibition of ACE2 and TMPRSS2 would reduce the spread of SARS-CoV-2 infection in a given COVID-19-(+) patient and this would allow the innate immune system and antivirals such as remdesivir to more effectively suppress early infection.

Due to its high pathogenicity and the lack of an effective treatment, live SARS-CoV-2 viruses must be handled under Biosafety Level 3 (BSL-3) conditions, which has hindered the development of vaccines and therapeutics. Pseudotyped viral particles are chimeric virions that consist of a surrogate viral core with a heterologous viral envelope protein at their surface. Such pseudoviruses are routinely used by many investigators for other highly pathogenic coronaviruses including SARS-CoV [42, 43] and MERS-CoV [44] to study viral entry, and develop assays for neutralizing antibodies and drug discoveries. For the current study, we have developed a pseudotyped SARS-CoV-2 virus which has a lentiviral core but with the SARS-CoV-2 spike protein on its envelope. The pseudoviruses infect human lung epithelial cells in an ACE2-dependent manner and confer the expression of a fluorescence protein ZsGreen in infected cells for imaging and quantification. The pseudoviruses can only accomplish a single infection cycle and are replication incompetent, thus require only BSL-2 level containment.

Our results suggest that MEK inhibitors, as a class, suppress host SARS-CoV-2 infectivity factors such as ACE2 and TMPRSS2, and that alone or in combination with remdesivir, there is innate immune system activity along with suppression of inflammatory cytokines and stimulation of Natural Killer cell activity. Our results support the further investigation of MEK inhibitors as a strategy to dampen early SARS-CoV-2 infection to allow host immunity as well as potentially antiviral agents to be more effective.

## Results

### MEK inhibitors reduce ACE2 expression in human cell lines

Based on prior literature that SARS coronavirus SPIKE protein through ACE2 can activate MAPK signaling [39], we hypothesized that MEK inhibitors (MEKi) may inhibit SARS-CoV-2 cellular effects. We used human tumor cell lines as well as normal human lung cells as a model to test effects of MEKi on ACE2 expression. We initially observed in H1975 human non-small cell lung cancer (NSCLC) cells that at doses below IC50, three different MEKi’s suppressed ACE2 protein expression (Figure 1A). VS-6766 (5 μM), a small molecule RAF/MEK inhibitor, MEKi Selumetinib (20 μM), or MEKi Trametinib (5 μM) all inhibited expression of ACE2 protein (as detected by PAB13444) with more subtle effects detected by another ACE2 antibody (CS4355) that recognizes glycosylated ACE2. We include results with the two commercially available antibodies we used to demonstrate that these antibodies did not always give concordant results. In this experiment the reduction in ACE2 was clearly demonstrated with the PAB13444 antibody. We observed that the cleaved active SP-domain of TMPRSS2 was increased by chloroquine or hydroxychloroquine and this was potentiated by the MEKi’s (Figure 1A). Inflammatory cytokine IL-6 was reduced by all 3 MEKi’s (RAF/MEKi VS-6766 showed the greatest reduction in this experiment) with no benefit from addition of chloroquine or hydroxychloroquine.

**Figure 1.**
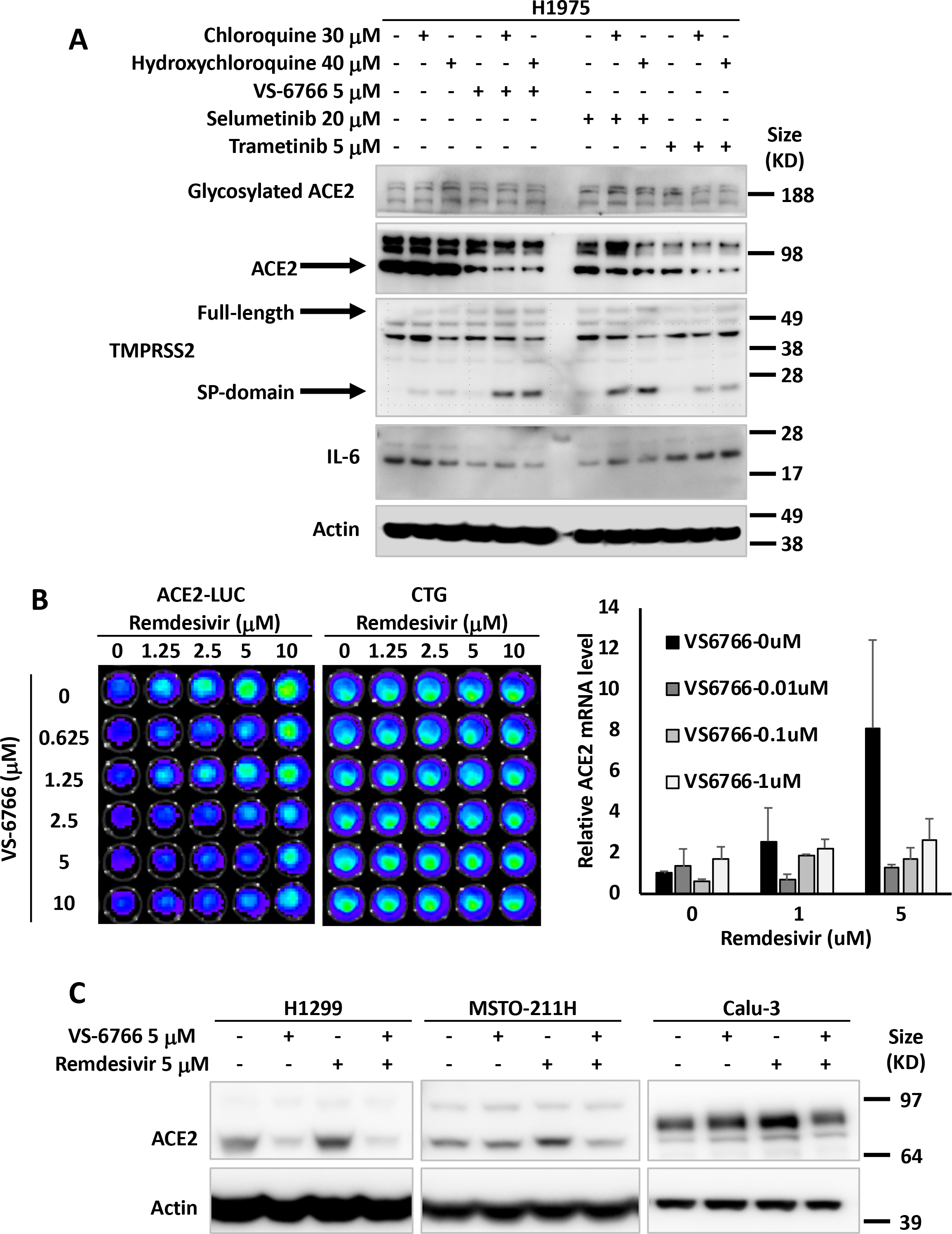
MEK inhibitors alone or in combination with remdesivir suppress ACE2 expression. The effects of chloroquine, hydroxychloroquine, remdesivir and MEK inhibitors (VS-6766, selumetinib and trametinib) on ACE2, TMPRSS2 and IL-6 in human lung and colon cells are shown. (A) H1975 human NSCLC cells were treated with the indicated drugs and doses for 48 hours. Glycosylated ACE2, ACE2, TMPRSS2 (full-length and Serine Protease-domain), IL-6 were probed with the cell signaling 4355, Abnova PAB13444, Sigma MABF2158, and Sigma SAB1408591 antibodies. β-Actin was probed with Sigma A5441 as a loading control. (B) Effect of remdesivir and VS-6766 on ACE2(−1119)-Luc reporter and cell viability. HCT116 human colorectal cancer cells were transiently transfected with ACE2(−1119)-Luc reporter for 24 hours followed by remdesivir and VS-6766 treatment for 24 hours at the indicated doses. First D-Luciferin was added to acquire ACE2-Luc reporter bioluminescence images (left panel) with the Xenogen IVIS system. After the bioluminescence signal decayed, CellTiter-Glo was added to acquire cell viability images (right panel; “CTG”) with the IVIS system. (C) Remdesivir alone increased ACE2 protein expression. H1299 human NSCLC cells, MSTO-211H human mesothelioma cells, and Calu-3 human NSCLC type II alveolar cells were treated with remdesivir and VS-6766 for 48 hours. ACE2 was probed with Abnova PAB13444 antibody, and β-Actin was probed with Sigma A5441 antibody as a loading control.

We introduced an ACE2-promoter luciferase-reporter (ACE2-luc) in HCT116 human tumor cells to investigate whether MEKi could inhibit expression of ACE2 from its endogenous promoter. Because of data that the anti-viral agent remdesivir could reduce hospitalization of COVID-19 infected patients [19], we investigated the effects of combining remdesivir on ACE2 expression. Surprisingly, we found that in HCT116 cells, ACE2-luc reporter expression was increased by remdesivir, and this was attenuated by the addition of VS-6766 at non-toxic doses (Figure 1B). Remdesivir also increased ACE2-Luc reporter activity in Calu-6 lung cancer cells at doses that did not reduce cell viability (Supplementary Figure 1A). The increase in ACE2-Luc reporter activity following remdesivir treatment of HCT116 cells was observed with three different ACE2-Luc promoter constructs containing 1119, 252, and 202 base pairs of the ACE2 promoter linked to firefly luciferase (Supplementary Figure 1B, C). VS-6766 attenuated at multiple doses with all 3 reporter constructs transfected into HCT116 cells (Supplementary Figure 1B, C). Similar results with induction of reporter activity after remdesivir treatment were observed and attenuated reporter activity after cotreatment with a second MEKi trametinib following ACE2-Luc reporter transfection in HCT116 cells (Supplementary Figure 1D, E). Similar results were obtained following reporter transfection and treatment with remdesivir and VS-6766 in an experiment performed by a different team member (Supplementary Figure 1F).

To extend the ACE2-Luc reporter studies, we evaluated the effects of remdesivir at 1 and 5 μM and VS-6766 (0.01, 0.1, and 1 μM) on ACE2 mRNA expression in HCT116 cells. We found a dose-dependent increase in ACE2 mRNA expression in HCT116 cells treated with remdesivir that was inhibited by addition of VS-6766 (Figure 1B). These results confirm that ACE2 mRNA expression was increased in remdesivir-treated HCT116 cells and this increase was attenuated at multiple doses of RAF/MEKi VS-6766.

To further evaluate the effects of MEKi plus remdesivir on ACE2 expression, we evaluated ACE2 protein expression in different human cell lines. We treated H1299, MST0211H, and Calu-3 cells with 5 μM remdesivir, 5 μM VS-6766 or the combination. ACE2 protein levels were reduced in all 3 cell lines when treated with VS-6766. Remdesivir increased ACE2 protein expression in all 3 cell lines and the combination of remdesivir plus VS-6766 led to reduced expression of ACE2 in all three cell lines. Thus, we found that several human cell lines increase ACE2 protein expression following treatment with remdesivir and this effect was inhibited by the addition of RAF/MEKi VS-6766 (Figure 1C).

Additional experiments were performed to test the effects of remdesivir and VS-6766 or other MEKi on ACE2, TMPRSS2, and IL-6 in tumor and normal cell lines. There was variability in the observed effects with some cell lines showing subtle induction of ACE2 protein expression by remdesivir or less robust suppression by MEKi. This was the case with HCT116 and Calu-6 cells (Supplementary Figure 2A). In HT-29 colorectal cancer cells and H1299 lung cancer cells, we observed that choloroquine and hydroxychloroquine increase expression of ACE2, TMPRSS2 and IL-6 (Supplementary Figure 2B). All these effects were suppressed by the 3 MEKi’s tested (VS-6766, selumetinib, and trametinib) in both HT-29 and H1299 cells (Supplementary Figure 2B). In BEAS-2B cells, there were subtle inductions of ACE2 expression after remdesivir and modest reductions in ACE2 expression after treatment by higher doses of VS-6766 (Supplementary Figure 3A). In H522 cells, there were little or no effects of remdesivir on ACE2 levels and minimal effects of VS-6766 on ACE2 expression at the highest doses used (Supplementary Figure 3B). in a NSCLC patient-derived cell line, we observed induction of ACE2 as well as TMPRSS2 after remdesivir treatment and these effects were suppressed by multiple doses of VS-6766 with especially notable effects on cleaved active TMPRSS2 (Supplementary Figure 3C). Further examples of variability in effects of hydroxychloroquine, remdesivir and VS-6766 are shown in Supplementary Figure 3D in normal and cancerous lung cells.

### Remdesivir and MEKi are nontoxic to normal and cancerous cells at doses that modulate SARS-CoV-2 infectivity factors, immune effects and cytokine levels

Given our interest in using MEKi treatment as a strategy to suppress SARS-CoV-2 infectivity either alone or in the presence of anti-viral agent remdesivir, we tested nontoxic doses of the drugs in the various experiments that are shown. Data in support of the lack of toxicity of the drugs are shown in Supplementary Figures 6-8.

### Recombinant SPIKE protein fragments can increase phospho-ERK and this is suppressed by MEK inhibitors

We tested whether SARS-CoV-2 SPIKE protein could increase MAPK and ERK signaling as was shown previously with SARS coronavirus [39]. We added fragments of recombinant SPIKE protein to human cells in culture and observed an increase in ACE2 expression (as detected by the SC390851 antibody) in Calu-3 cells (Figure 2A). ACE2 expression was suppressed by MEKi inhibitor treatment of Calu-3 non-small cell lung cancer cells (Figure 2A). We further observed that in HT-29 colon cancer cells and BEAS-2B human bronchial airway epithelial cells, pERK was increased by SPIKE protein, while the addition of RAF/MEKi VS-6766 inhibited expression of both pERK and ACE2 (Figure 2B). Total ERK levels were unchanged in these cells (Figure 2B). The recombinant SPIKE fragments did not appear to increase pERK in the experiment in Calu-3 cells (Figure 2A).

**Figure 2.**
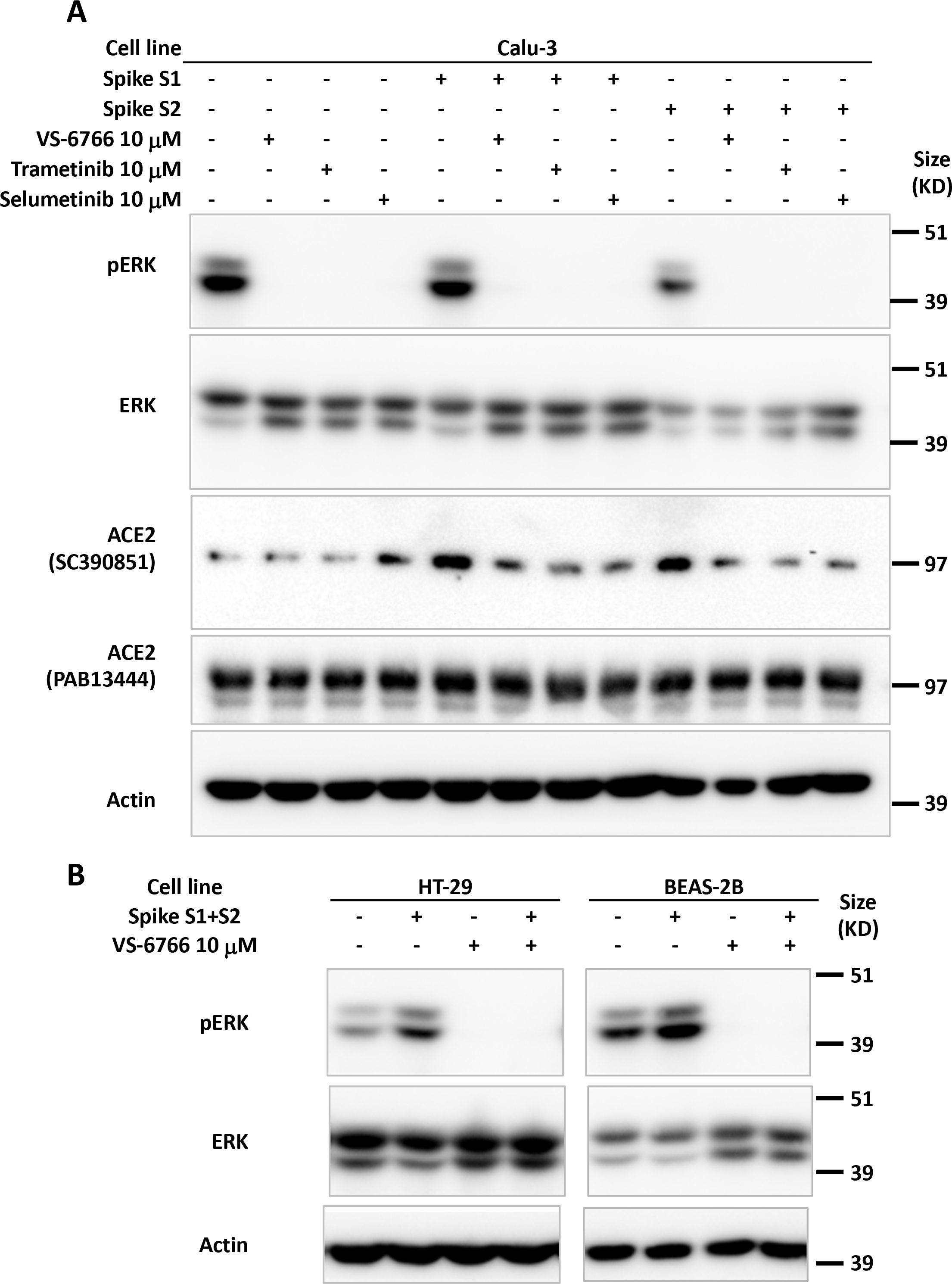
Increase in pERK by recombinant SPIKE protein is suppressed by MEK inhibition. (A) Calu-3 human NSCLC type II alveolar cells were pretreated with 10 μM MEK inhibitor (VS-6766, trametinib or selumetinib) for 1 hr before incubation with 4.7 μM recombinant SARS-CoV-2 spike protein subunit 1 (S1, 0.118 μg/mL) or subunit 2 (S2, 0.275 μg/mL) for 2 hr. ACE2 protein detected with sc-390851 was increased following cell incubation with S1 and S2. All three MEK inhibitors suppressed increased ACE2. (B) HT-29 human colorectal cancer cells and BEAS-2B normal human bronchial epithelial cells were pretreated with 10 μM RAF/MEK inhibitor VS-6766 for 1 hr before incubation with 4.7 μM recombinant SARS-CoV-2 spike protein subunit 1 (S1, 0.118 μg/mL) and subunit 2 (S2, 0.275 μg/mL) for 2 hr. Recombinant SARS-CoV-2 spike protein subunits increased pERK expression which was completely abrogated by VS-6766 treatment.

### Suppression of ACE2 protein expression in correlation with reduction of pERK after MEKi treatment of human cells

To further investigate the correlation between MAPK-pERK activation and ACE2 expression we investigated the effects of remdesivir and VS-6766 on ACE2 expression and pERK expression in several human cell lines (Figure 3). We observed a correlation between pERK expression and ACE2 protein expression under a number of different experimental conditions including treatment of cells with remdesivir, VS-6766, or the combination, as well as serum deprivation and restimulation conditions (Figure 3). For example, under standard cell culture conditions, remdesivir increased pERK and ACE2 levels in HCT116, H1975, and BEAS-2B cells while treatment with VS-6766 alone or in combination with remdesivir suppressed both pERK and ACE2 levels (Figure 3A). In MRC-5 normal human lung fibroblast cells, we observed an increase in pERK following remdesivir treatment under normal culture conditions or following serum stimulation of serum-deprived cells (Figure 3B). We used serum deprivation and restimulation as a strategy to modulate MAPK signaling to further explore the effects of remdesivir and MEKi. Both pERK and ACE2 levels were suppressed by treatment of MRC-5 cells with VS-6766 alone or in combination with remdesivir in serum deprived or serum-deprived and subsequently serum-stimulated MRC-5 cells (Figure 3B). An increase in pERK was observed in H460 and A549 lung cancer cells following remdesivir treatment and this was suppressed by VS-6766 alone or in combination with remdesivir (Figure 3C, D). In Calu-3 cells, we observed an increase in pERK that was suppressed by addition of VS-6766 either following serum deprivation or serum stimulation of serum-deprived cells (Figure 3E, right panel) while at later time points ACE2 expression was inhibited by VS-6766 plus remdesivir (Figure 3E, left panels). Thus, pERK was increased by remdesivir treatment under multiple experimental conditions, and this was correlated with expression of ACE2 (Figure 3). With VS-6766, both pERK and ACE were suppressed, including when VS-6766 was combined with remdesivir (Figure 3).

**Figure 3.**
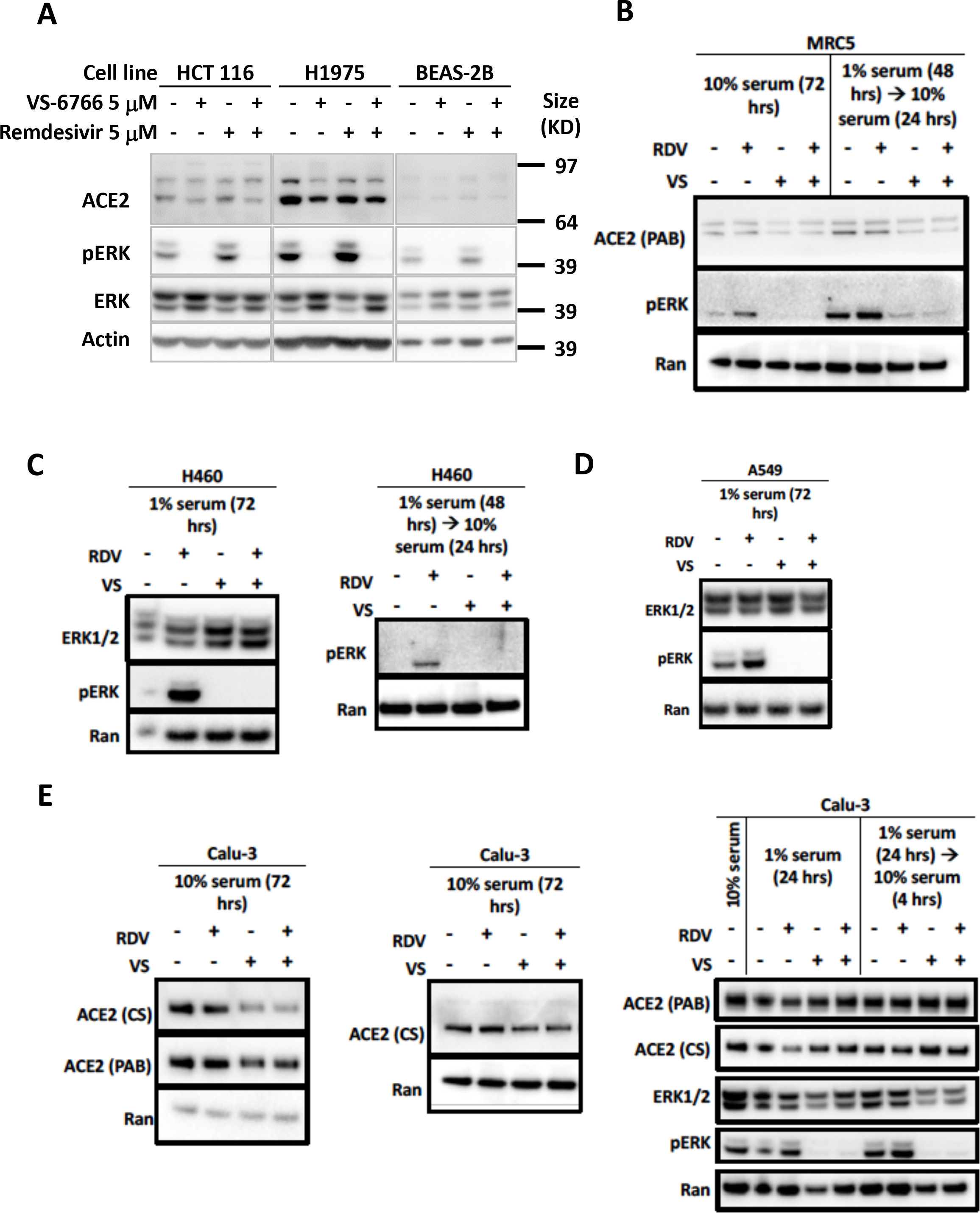
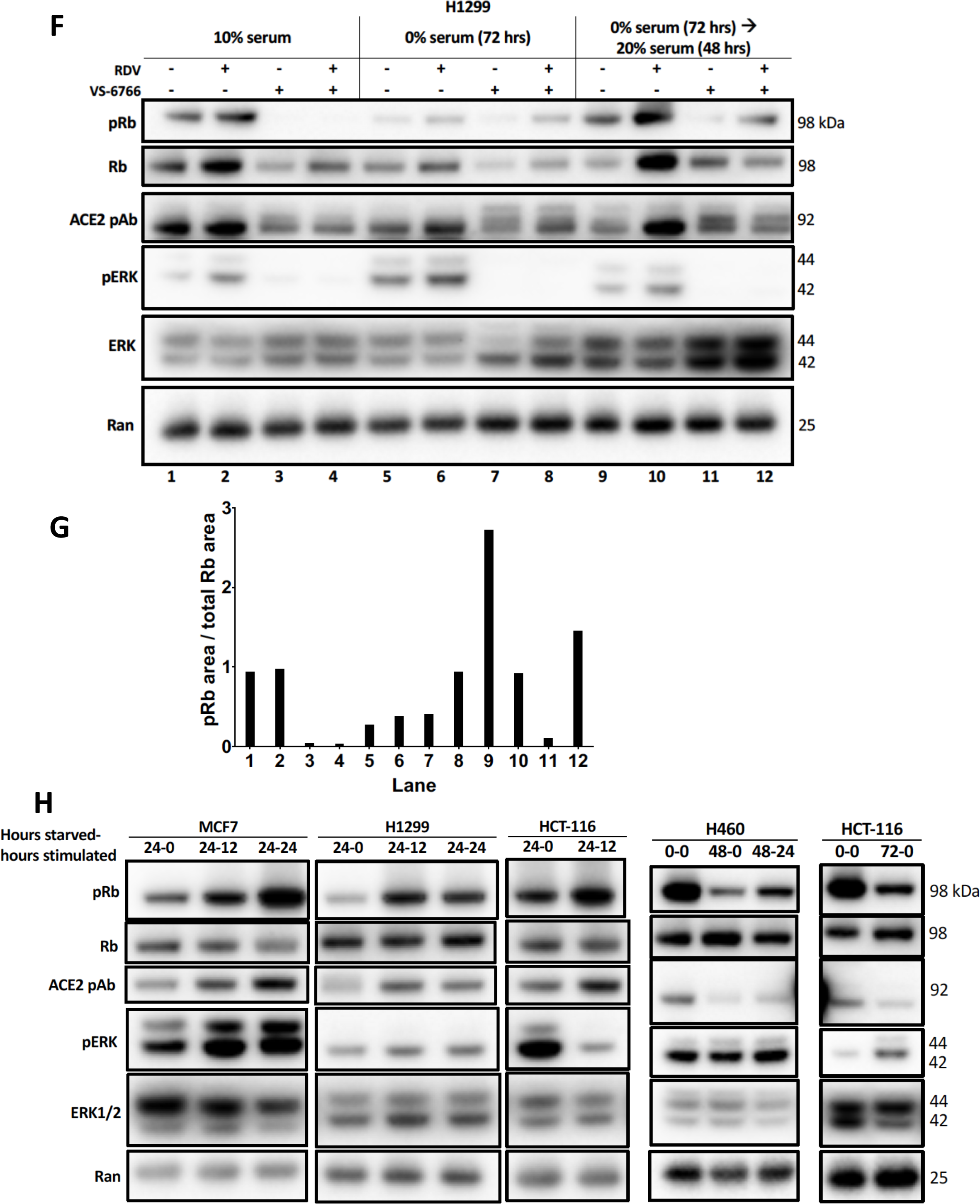
Reduced phospho-ERK and ACE2 expression by the combination of MEK inhibitor and remdesivir. (A) Effects of 5 μM remdesivir (RDV), RAF/MEK inhibitor VS-6766, or the combination on ACE2, ERK1/2, and pERK protein expression. HCT116 human colorectal cancer cells, H1975 human NSCLC cells and BEAS-2B normal human bronchial epithelial cells were treated with 5 μM VS-6766 or/and 5 μM Remdesivir for 48 hr. Remdesivir alone increased pERK protein expression in all three cell lines, which was completely depleted by VS-6766 treatment. Effects of 5 μM RDV, VS-6766, or the combination on ACE2, ERK1/2, and pERK protein expression in (B) normal human lung cells and (C-E) human lung cancer cells are shown (B-E left 2 panels). Serum-deprived cells were plated and cultured in medium containing 1% FBS for the indicated amount of time. For serum-stimulated cells, FBS was added to a final concentration of 10% for 24 hours (E, right panel). Cells were plated in 10% serum for 16 hours, then media was removed and replaced with 1% FBS for serum-deprived cells. For serum-stimulated cells, FBS was added to a final concentration of 10% for 4 hours. ACE2 (PAB), ACE2 (CS), ERK1/2, and pERK were probed with Abnova PAB13444, Cell Signaling 4355, Cell Signaling 9102, and Cell Signaling 4370 antibodies. Ran was probed with BD Biosciences 610341 antibody as a loading control. (F) Modulation of pRb and ACE2 expression in H1299 cells with serum starvation, stimulation, and MEKi treatment. Control cells were grown with the normal 10% serum throughout the treatment. Serum starved cells were plated in 10% serum and incubated for 16 hours, then grown in media containing 0% serum for 72 hours. Serum stimulated cells were similarly starved for 72 hours, then stimulated with media containing 20% serum for 48 hours. All drug treated cells received 5 mM RDV, VS-6766, or the combination 48 hours before harvesting. (G) pRb as a fraction of total Rb was quantified in ImageJ. (H) Modulation of ACE2 expression with serum starvation and stimulation. Four different cell lines (2 lung, 1 colorectal, and 1 breast cancer) were grown in 10% serum for 16 hours then were starved in 0% FBS for 0 (no starvation), 24, 48, or 72 hours. Cells were stimulated with 20% FBS and harvested either 0 (no stimulation), 12, or 24 hours later. An increase in pRb relative to total Rb was seen upon stimulation and this correlated with ACE2 levels. pERK correlation with ACE2 was heterogeneous, with a correlation seen in MCF7 cells but not H1299 or HCT-116 cells (three left-most panels). pRb relative to total Rb decreased upon starvation and increased upon stimulation, which correlated with ACE2 expression (H460 and HCT-116 in right-most panels).

### Correlation between pERK, pRb and ACE2 in serum-deprived and -restimulated cells with and without remdesivir and MEK inhibition

We performed serum deprivation and re-stimulation to further test correlations between the status of activation of the MAPK pathway, proliferation state, and ACE2 expression. The data shows a correlation between phospho-Rb, and ACE2 expression (Figure 3F-H). This is in addition to further replication of the effects of remdesivir and MEKi on ACE2 in these experiments where we observe correlations between pERK, pRb, and ACE2 (Figure 3F). Serum-starved cells showed a decrease in pRb levels compared to the 10% FBS control, which correlated with a decrease in ACE2 (Figure 3F, lanes 1 and 5). An increase in pRb/total Rb was observed under serum stimulation, but with no increase in ACE2 (Figure 3F, compare lanes 5 and 9). VS-6766 and the combination treatment decreased pERK, which under normal conditions correlated with a decrease in pRb relative to total Rb and a reduction in ACE2 levels. Remdesivir increased ACE2 expression under all conditions, most significantly under stimulation (Figure 3F). An increase in pRb relative to total Rb was observed upon serum stimulation and this correlated with ACE2 levels (Figure 3H). pERK correlation with ACE2 was heterogeneous, with a correlation seen in MCF7 cells but not H1299 or HCT-116 cells (Figure 3H). In H460 and HCT116 cells, pRb relative to total Rb decreased upon starvation and increased upon stimulation, which correlated with ACE2 expression (Figure 3H, right panels). Thus, the results in Figure 3F show the triple correlation between pERK/pRb/ACE2 after MEKi and/or remdesivir treatment, and shows correlation of pRb/ACE2 when evaluating untreated cells before and after starvation. We quantified pRb and show as a fraction of total Rb (Figure 3G). Figure 3H similarly shows that under conditions of starvation/stimulation, there is a correlation between pRb and ACE2.

### MEK inhibitors stimulate Natural Killer cell (but not T-cell) activity against target cells

Since we observed that MEKi can inhibit ACE2 expression to potentially reduce SARS-CoV-2 infectivity, we further investigated effects of MEKi as well as remdesivir on Natural Killer (NK) cell activity against target cells. NK cells serve as a natural defense against transformed and virally infected cells including through cytotoxic ligands such as TRAIL. Green fluorescent tumor cells were co-cultured with NK-92 cells at a 1:1 effector target cell ratio (E:T) for 24 hours after treatment with 1 μM of the indicated drugs (Figure 4). Our results reveal that MEKi can stimulate target cell killing by NK cells (Figure 4). Similarly designed experiments did not reveal stimulation of T-cell killing of target cells by various MEKi (Supplementary Figure 9). We performed the immune cell killing assays under conditions where MEKi were generally non-toxic to target cells or immune cells (Supplementary Figures 10-12).

**Figure 4.**
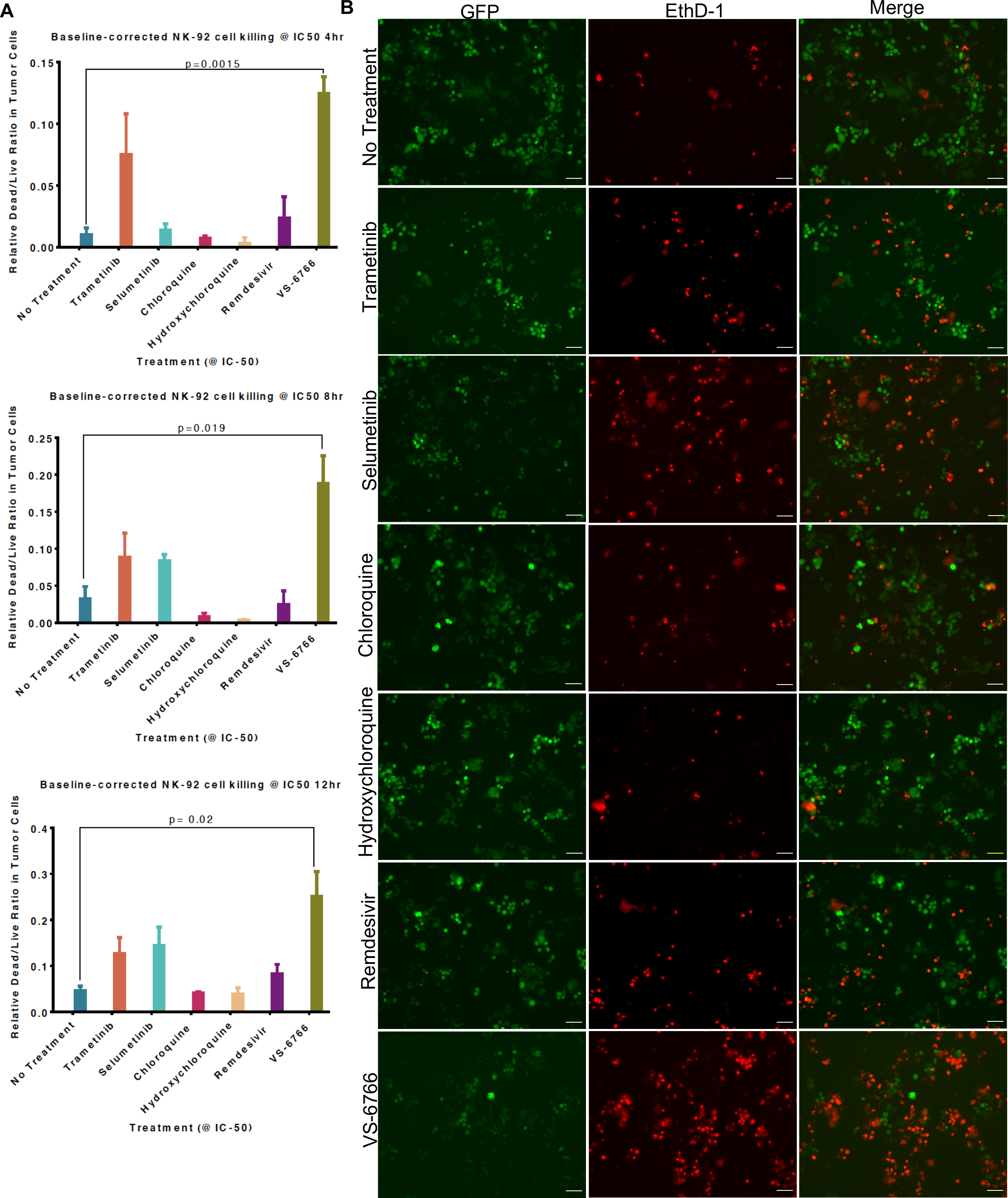
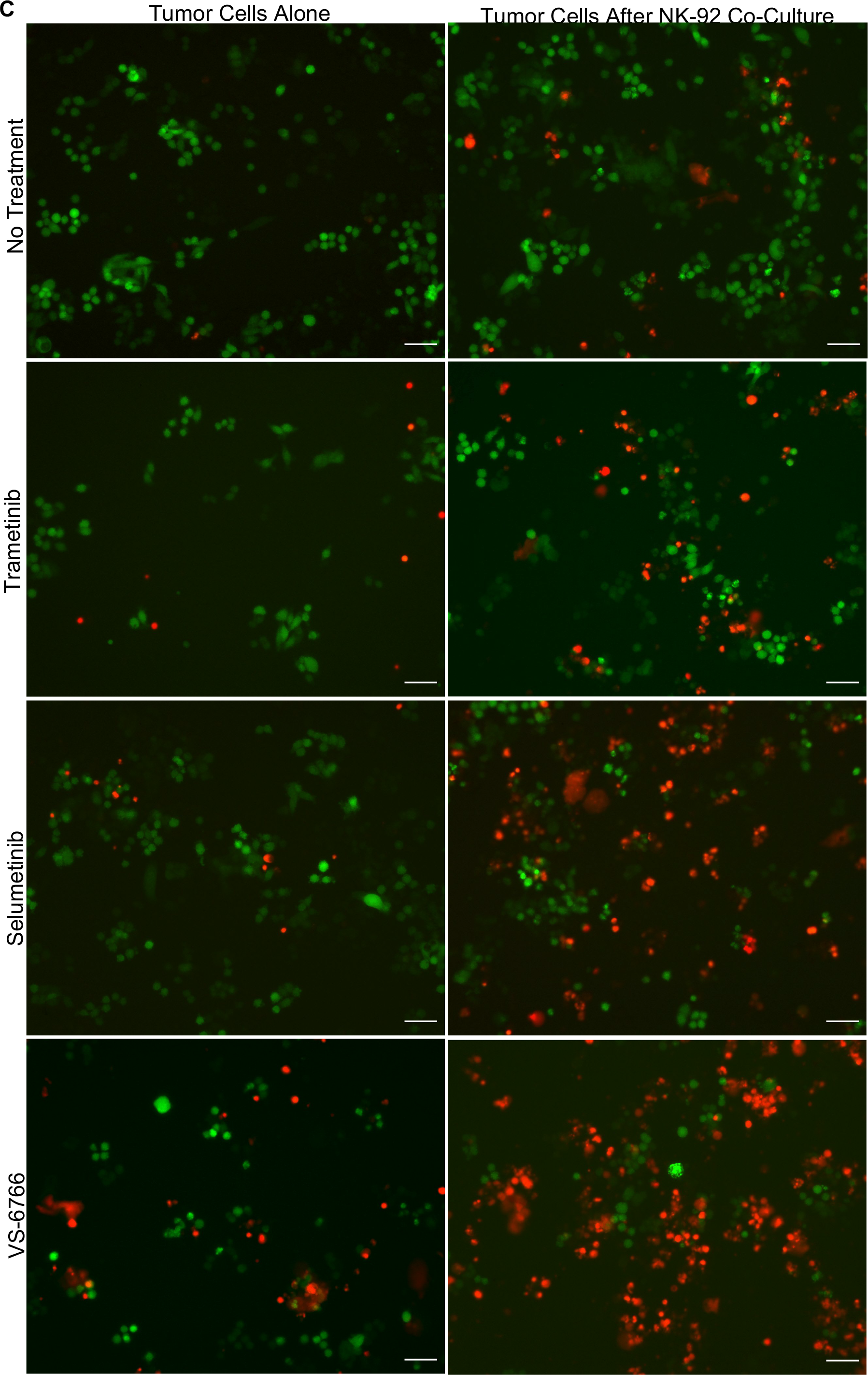
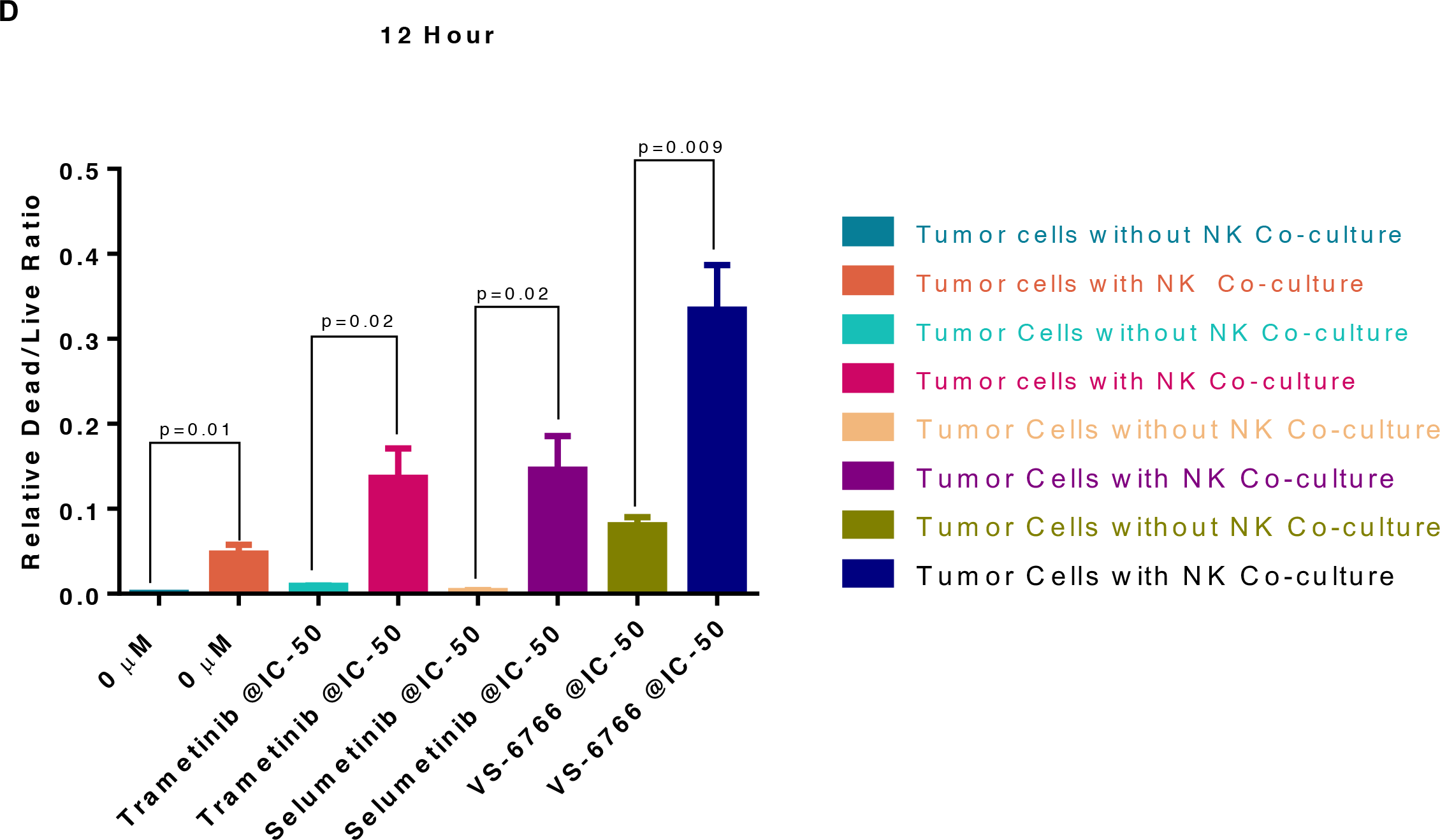
Stimulation of Natural Killer activity by MEK inhibitor treatment at IC-50 doses. Green fluorescent SW480 tumor cells were co-cultured with NK-92 natural killer cells at a 1:1 effector target cell ratio (E:T) for the indicated timepoints and imaged by fluorescence microscopy. Cells were treated with the indicated drug at IC-50 doses and drugs were added at the same time as the NK cells. (A) Quantification of dead/live ratio after 4, 8 and 12 hours of treatment by drugs as indicated. P values are displayed on graph and were calculated using unpaired t tests. (B) Fluorescence microscopy of GFP+ SW480 tumor cells before and after 12 hours of indicated treatment conditions. Ethidium homodimer was used to visualize dead cells. 10x magnification. Scale bars indicate 100 μM. (C) Images showing GFP+ tumor target cell cytotoxic effects of MEK inhibitors alone or in addition to NK-92 cells after 12 hours of indicated treatment conditions. Ethidium homodimer was used to visualize dead cells. 10x magnification. Scale bars indicate 100 μM. (D) Quantification of tumor target cell cytotoxic effects of MEK inhibitors alone or in addition to NK-92 cells. P values are displayed on graph and were calculated using unpaired t tests.

### Increased cytokine expression in plasma from COVID-19-(+) patient plasma samples

As a pilot study, we set up a screening cytokine array to evaluate cytokine levels in nine COVID-19-(+) patient plasma versus eleven normal healthy volunteer samples. Patient characteristics, symptoms, diagnoses, and interventions are shown in Tables 1-4. Cytokine levels observed in plasma samples from specific patients are as indicated in Table 5. The cytokine profiling results reveal significantly increased levels of M-CSF (P=0.001), IL-6 (P=0.042), IL-1RA (P=0.028), IP-10 (P=0.02), IFNα2 (P=0.013) and TNF-a (P=0.0072) and trends towards significance for MCP-1 (P=0.055), IFN-γ (P=0.1), IL-2 (P=0.15), IL-7 (P=0.17), G-CSF (P=0.11) in COVID-19-(+) plasma samples (Figure 5A). IP-10 is also known as CXCL10 or interferon gamma-induced protein 10 that is produced by monocytes, endothelial cells and fibroblasts and serves as a chemoattractant for immune cells. No appreciable changes were observed in IL-12p40, IL-18, or IL-1A in COVID-19-(+) plasma samples versus controls. The highest levels of cytokines G-CSF, MCP-1, IFN-γ, IL-1RA, IL-6, IP-10, M-CSF, IL-2, IL-1A, and TNF-alpha were observed in the sickest ICU-admitted COVID-19-(+) patients #2 and 103. For the individual COVID-19-(+) patients the cytokine levels are listed in Table 5. The cytokine values in patient plasma are shown individually graphically for each patient in Supplementary Figure 4.

**Figure 5.**
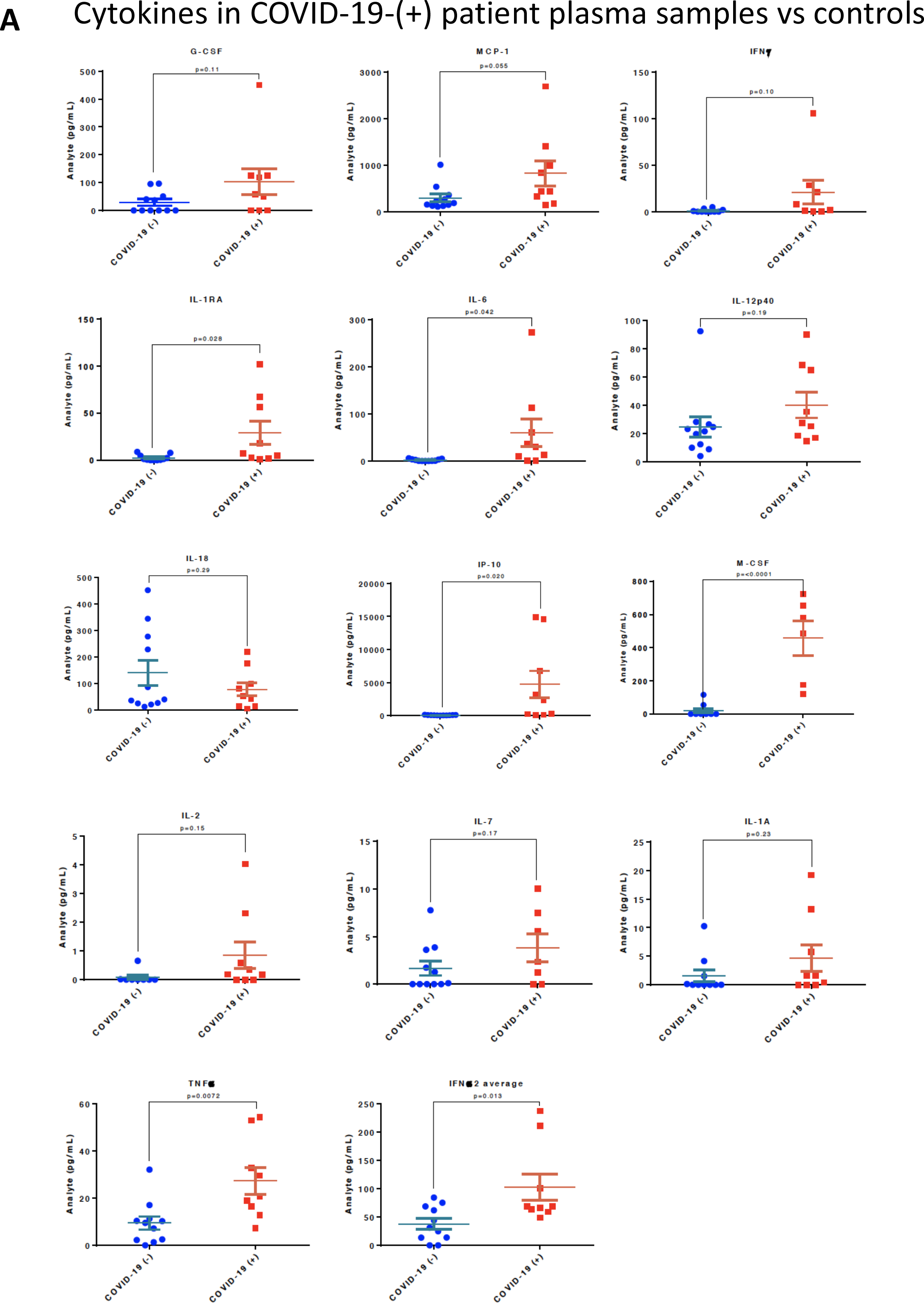
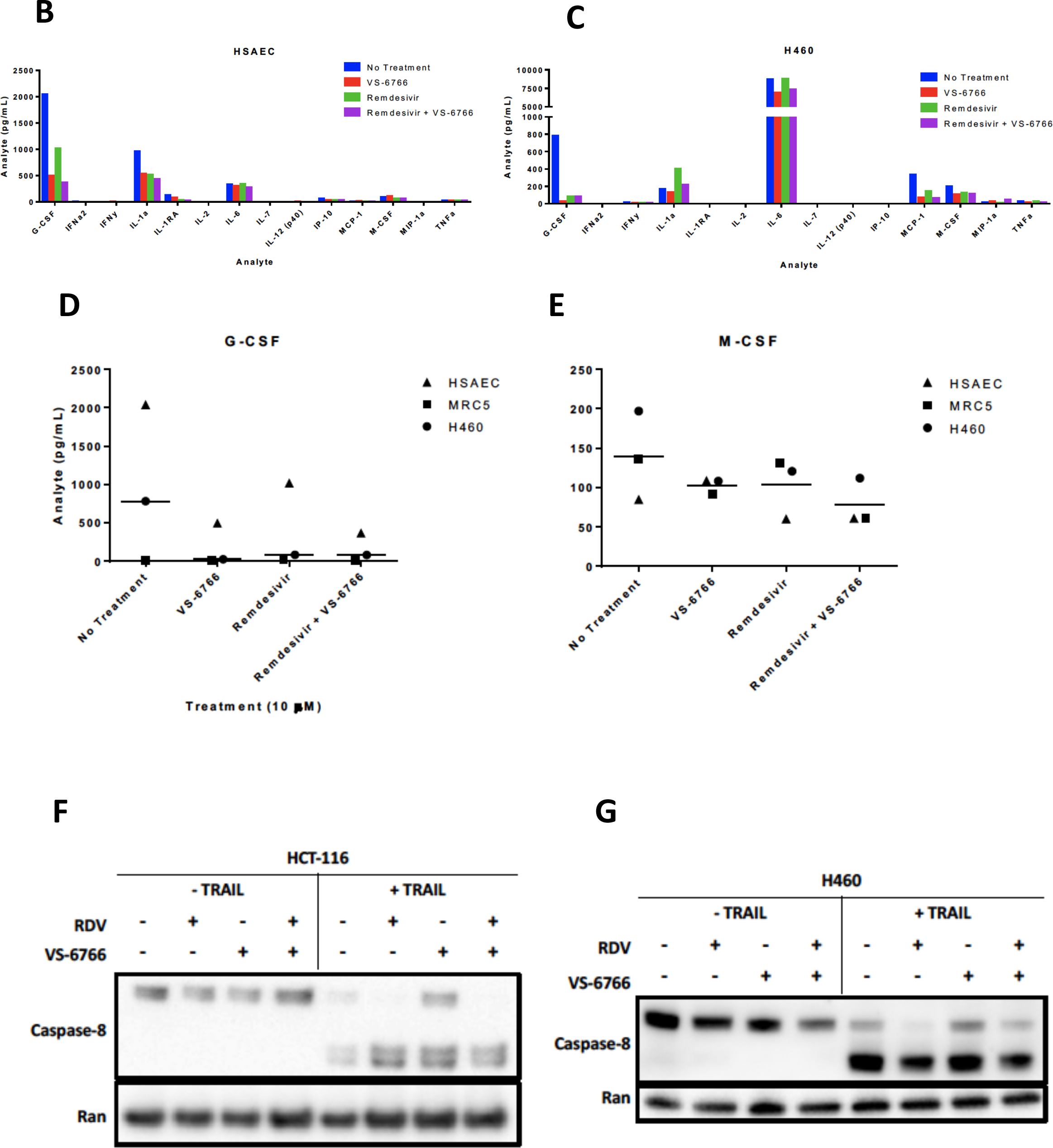
Increased cytokines in COVID-(+) patient plasma and demonstration that VS-6766 plus remdesivir reduces cytokine secretion while allowing TRAIL mediated killing of target cells. (A) Cytokine levels in COVID-19-(+) patient plasma versus normal patient controls. P-values are as indicated for each cytokine measurement. For the COVID-19-(+) patients, the cytokine levels for individual patients are shown in Supplementary Figure 4. (B-E). Cytokine profiles of normal and cancerous lung cells. HSAEC, MRC-5, and H460 cells were treated for 48 hours with the indicated drugs. (B) Cytokine profiles of HSAEC after treatment. (C) Cytokine profiles of H460 cells after treatment. (D) Comparison of G-CSF levels in cell types between treatment conditions. (E) Comparison of M-CSF levels in cell types between treatment conditions. Effects of remdesivir (5 μM), VS-6766 (5 μM), or the combination for 24 hours on (50 ng/mL; 4 additional hours) TRAIL-mediated activation of cleaved caspase 8 in HCT116 colorectal cancer (F) or H460 lung cancer cells (G).

**Table 1.** Patient History

| Record ID | Age | Sex | Race | No Health Insurance | Private Insurance | Medicare | Medicaid | Smoking | Chronic lung disease | Chronic kidney disease | Chronic neurologic disease | Heart disease | High blood pressure | Diabetes | Auto-immune disease | HIV/AIDS | Active cancer | Previous stroke | Obesity |
| --- | --- | --- | --- | --- | --- | --- | --- | --- | --- | --- | --- | --- | --- | --- | --- | --- | --- | --- | --- |
| 1 | 43 | Male | Black | No | No | Yes | No | Quit smoking | No | No | No | Yes | No | Yes | No | No | No | No | No |
| 2 | 67 | Male | Black | No | Yes | Yes | Yes | No history | No | No | No | No | Yes | Yes | No | No | No | No | No |
| 4 | 64 | Male | Other | No | No | No | Yes | No history | No | Yes | No | Yes | Yes | Yes | No | No | No | Yes | No |
| 5 | 73 | Male | White | No | No | Yes | Yes | Quit smoking | Yes | No | No | Yes | Yes | No | No | No | No | Yes | No |
| 6 | 50 | Male | Other | No | No | No | Yes | No history | Yes | No | No | No | No | Yes | No | No | No | No | No |
| 8 | 57 | Male | Other | No | No | No | Yes | No history | No | No | No | No | Yes | No | No | No | No | No | No |
| 35 | 33 | Male | Other | No | Yes | No | No | No history | No | No | No | No | No | No | No | No | No | No | No |
| 103 | 73 | Male | White | No | No | Yes | Yes | Quit smoking | Yes | No | No | No | Yes | No | No | No | No | No | Yes |
| 122 | 51 | Male | Unknown | No | No | No | Yes | No history | No | No | No | No | Yes | No | No | No | No | No | No |

**Table 2.** Patient Symptoms

| Record ID | Breathing Difficulty or Shortness of Breath | Fever | Cough | Change in taste or smell | Rash | GI symptoms | Neuro symptoms | Sore throat | Chest pain | Systolic BP | Diastolic BP | Pulse | RR | Temperature | Is the patient requiring supplemental oxygen? | How much oxygen is the patient on? | What is the patient's pulse oximetry reading? |
| --- | --- | --- | --- | --- | --- | --- | --- | --- | --- | --- | --- | --- | --- | --- | --- | --- | --- |
| 1 | No | No | No | No | No | No | No | No | No | 140 | 93 | 73 | 18 | 99.5 | No |  |  |
| 2 | Yes | Yes | Yes | Yes | No | No | No | No | No | 167 | 75 | 106 | 22 | 98.1 | No |  |  |
| 4 | No | Yes | Yes | No | No | Yes | No | Yes | Yes | 205 | 109 | 78 | 22 | 98.9 | No |  |  |
| 5 | Yes | No | No | No | No | No | No | No | No | 81 | 58 | 66 | 22 | 97.4 | Yes | 4 - 6 L | 93 |
| 6 | Yes | Yes | Yes | No | No | No | No | No | Yes | 127 | 74 | 93 | 24 | 98.4 | No |  |  |
| 8 | Yes | Yes | No | No | No | No | No | No | No | 177 | 98 | 89 | 29 | 99.9 | Yes | >6 L | 91 |
| 35 | No | No | No | No | No | Yes | Yes | No | No | 176 | 111 | 82 | 18 | 98 | No |  |  |
| 103 | Yes | Yes | Yes | No | No | Yes | Yes | No | No | 120 | 82 | 108 | 18 | 96.5 | Yes | 4 - 6 L | 99 |
| 122 | Yes | Yes | Yes | No | No | Yes | No | Yes | No | 133 | 74 | 93 | 18 | 102.7 | Yes | >6 L | 88 |

**Table 3.** Emergency Department Diagnosis

| Record ID | ED Disposition | COVID-19 Diagnosis | Pneumonia Diagnosis | Other Diagnosis | COVID-19 PCR Testing | Chest X-ray |
| --- | --- | --- | --- | --- | --- | --- |
| 1 | Discharged - home | Yes | No | No | Positive | Normal |
| 2 | Admission - ICU or Step-down | Yes | Yes | No | Positive | Abnormal |
| 4 | Admission - floor | Yes | Yes | No | Positive | Abnormal |
| 5 | Admission - floor | Yes | No | No | Positive | Normal |
| 6 | Admission - floor | Yes | Yes | No | Positive | Abnormal |
| 8 | Admission - ICU or Step-down | Yes | Yes | No | Positive | Abnormal |
| 35 | Admission - ICU or Step-down | Yes | No | Yes | Positive | Normal |
| 103 | Admission - ICU or Step-down | Yes | Yes | Yes | Positive | Abnormal |
| 122 | Admission - ICU or Step-down | Yes | No | No | Positive | Abnormal |

**Table 4.** Interventions and Medications in the Emergency Department

| Record ID | Supplemental oxygen, < 6L/min | Supplemental oxygen, > 6L/min | BiPAP/CPAP | Incubation | Ibuprofen | Acetaminophen | Bronchodilators | Steroids | Azithromycin | Hydroxychloroquine | Antibiotic, other | Other |
| --- | --- | --- | --- | --- | --- | --- | --- | --- | --- | --- | --- | --- |
| 1 | No | No | No | No | No | No | No | No | No | No | No | No |
| 2 | No | Yes | No | No | No | Yes | Yes | No | No | No | Yes | No |
| 4 | No | No | No | No | No | No | No | No | No | No | No | No |
| 5 | Yes | Yes | No | No | No | No | No | No | Yes | No | Yes | No |
| 6 | Yes | No | No | No | No | No | Yes | No | No | No | No | No |
| 8 | No | Yes | No | No | No | Yes | No | No | No | No | No | No |
| 35 | No | No | No | No | No | Yes | No | No | No | No | No | Yes |
| 103 | Yes | No | Yes | No | No | No | No | No | No | No | No | Yes |
| 122 | Yes | No | No | No | No | No | No | No | No | No | No | Yes |

**Table 5.** COVID(+) Patient Analyte Values (pg/ml_)

| | G-CSF | IFNα2 | IFNγ | IL-1α | IL-1RA | IL-2 | IL-6 | IL-7 | IL-12 (p40) | IL-18 | IP-10 | MCP-1 | M-CSF | TNFα |
| --- | --- | --- | --- | --- | --- | --- | --- | --- | --- | --- | --- | --- | --- | --- |
| Patient 1 | 0.0 | 59.4 | 0.0 | 0.4 | 1.4 | 0.2 | 0.2 | 0.0 | 14.5 | 3.9 | 51.7 | 182.2 | 0.0 | 12.9 |
| Patient 2 | 450.5 | 211.3 | 105.5 | 19.2 | 101.9 | 4.0 | 273.5 | 5.5 | 64.9 | 41.6 | 14923.0 | 2690.0 | 581.6 | 54.5 |
| Patient 4 | 126.1 | 66.0 | 21.1 | 5.8 | 7.6 | 0.2 | 35.6 | 10.0 | 35.4 | 219.4 | 14632.0 | 991.4 | 176.4 | 29.7 |
| Patient 5 | 50.5 | 68.5 | 1.8 | 1.7 | 17.8 | 0.6 | 12.5 | 0.0 | 27.4 | 12.4 | 247.2 | 323.4 | 0.0 | 19.0 |
| Patient 6 | 119.0 | 69.3 | 1.0 | 1.6 | 2.1 | 0.4 | 0.3 | 2.4 | 18.4 | 13.4 | 138.0 | 149.6 | 0.0 | 7.3 |
| Patient 8 | 0.0 | 63.6 | 8.2 | 0.0 | 5.1 | 0.0 | 29.8 | 0.0 | 25.2 | 99.0 | 2381.0 | 435.3 | 121.8 | 16.4 |
| Patient 35 | 0.0 | 49.1 | 0.0 | 0.0 | 2.5 | 0.0 | 10.2 | 0.0 | 17.0 | 51.4 | 293.2 | 1400.0 | 652.9 | 32.7 |
| Patient 103 | 58.5 | 237.6 | 28.4 | 13.2 | 67.5 | 2.3 | 60.9 | 7.5 | 90.2 | 175.9 | 3159.0 | 830.7 | 724.7 | 53.0 |
| Patient 122 | 125.5 | 100.7 | 0.0 | 0.0 | 56.6 | 0.0 | 113.2 | 1.2 | 68.6 | 79.6 | 6815.0 | 430.4 | 487.9 | 20.7 |

### MEK inhibition is associated with reduced cytokine secretion by human cells

To further evaluate the potential use of MEKi to suppress SARS-CoV-2 infectivity and disease severity, we evaluated the impact of VS-6766 on cytokine release by various human cell lines in culture. Using a Luminex-200 multiplexed cytokine array read-out, we observed reduced cytokine release following cell treatment by the MEKi either used alone or in combination with remdesivir (Figure 5B-E). Cytokines that were suppressed include G-CSF, M-CSF, IL-1RA, MCP-1, IL-1a.

### MEK inhibition alone or with remdesivir does not suppress TRAIL-mediated killing of target cells

The TNF-related apoptosis-inducing ligand (TRAIL) is involved in killing transformed cells and virally-infected cells, with little inflammatory response. It is therefore important that any therapeutic agent being tested to control COVID-19 be monitored for effects on TRAIL-mediated killing of target cells. Having shown that MEKi’s can stimulate killing of target cells by NK cells (Figure 4), we further tested whether MEKi or remdesivir could suppress TRAIL-mediated killing of target cells. Our results reveal that RAF/MEKi VS-6766 either alone or in combination with remdesivir does not inhibit TRAIL-mediated killing of target cells (Figure 5F, G). Additional experiments confirmed these results and extended the findings to the other MEKi (Supplementary Figure 6).

### SARS-CoV-2 pseudovirus that expresses SPIKE protein variants on the envelope of a lentiviral core, infection of human airway epithelial cells or lung cancer cells, and demonstration of MEKi attenuation of infectivity

We developed a SARS-CoV-2 pseudovirus model system to investigate further our findings that MEKi may attenuate virus infectivity. We generated a pseudotyped SARS-CoV-2 virus which has a lentiviral core but with the SARS-CoV-2 D614 or G614 spike (S) protein on its envelope and used VSV-G lentivirus as a negative control (Figure 6A). Both SPIKE protein variants were expressed by the lentivirus (Figure 6A). The results suggest more G614 than D614 S protein was present on each viral particle. In order to establish an experimental model system for SARS-Cov-2 infection of human cells, we incubated SARS-CoV-2-S pseudovirus or control with BEAS-2B human bronchial airway epithelial cells or Calu-3 lung cancer cells. We observed GFP expression by fluorescence microscopy or by flow cytometry that is indicative of cellular infection (Figures 6B-E). In order to determine the effect of MEKi on infectivity by the SARS-CoV-2-S pseudoviruses, we pretreated Calu-3 cells with trametinib for 48 hrs and then infected the cells with the pseudoviruses and continued incubation in the presence of the MEKi (Figure 6F). This initial experiment was conducted in lung cancer cells which was not ideal and also under conditions where there was some cell loss and so the numbers of cells analyzed was more limited after trametinib treatment. We then chose to perform a second experiment using more normal lung epithelial cells and used shorter incubation times with the MEKi and analyzed at an earlier time point after pseudovirus infection. This subsequent experiment was performed in BEAS-2B normal lung airway epithelial cells and it showed inhibition of pseudovirus infection by various MEKi under conditions where most of the cells were surviving (Figure 6G). The results demonstrate inhibition of pseudovirus infectivity of human airway epithelial cells by MEKi.

**Figure 6.**
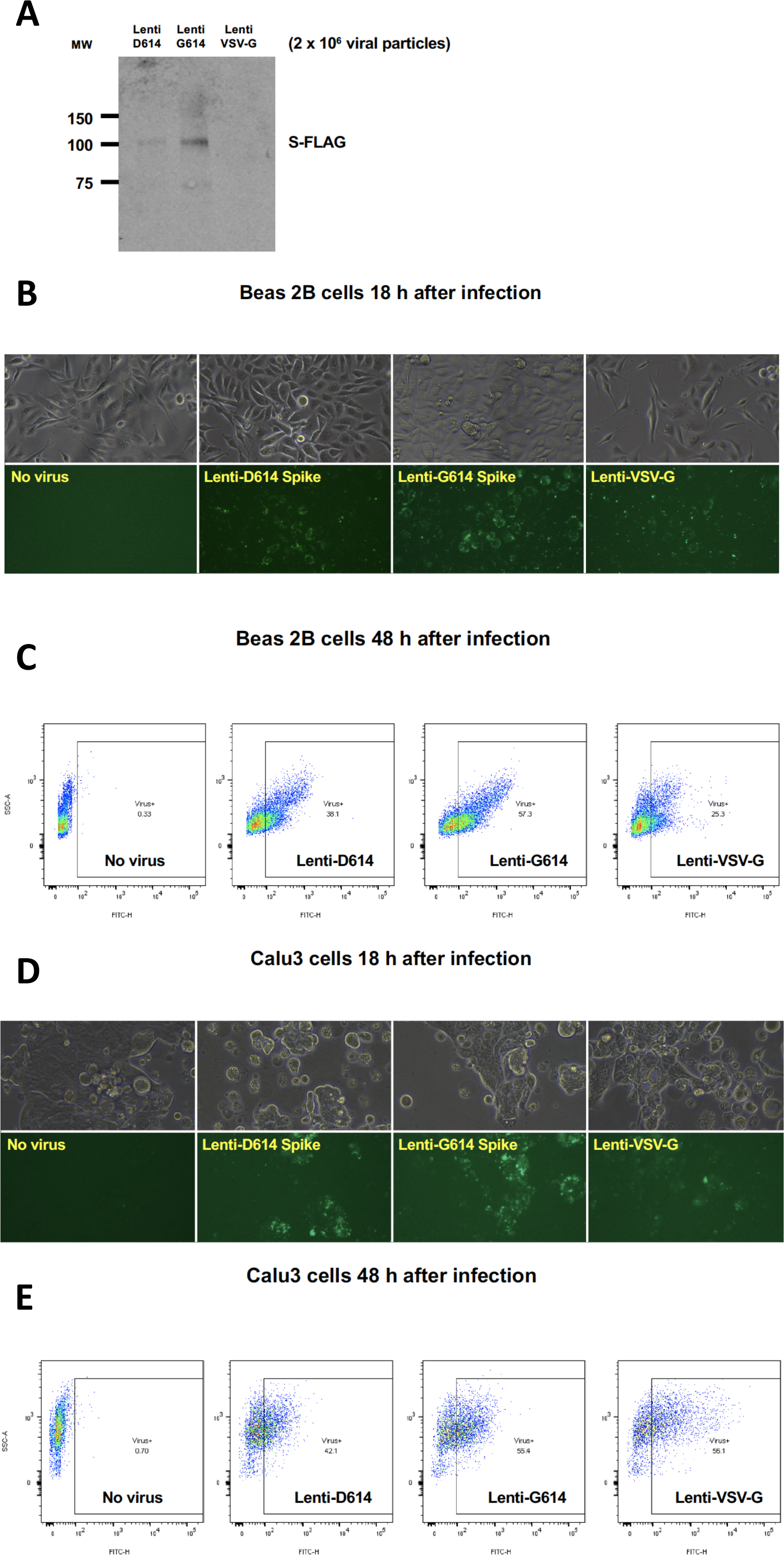
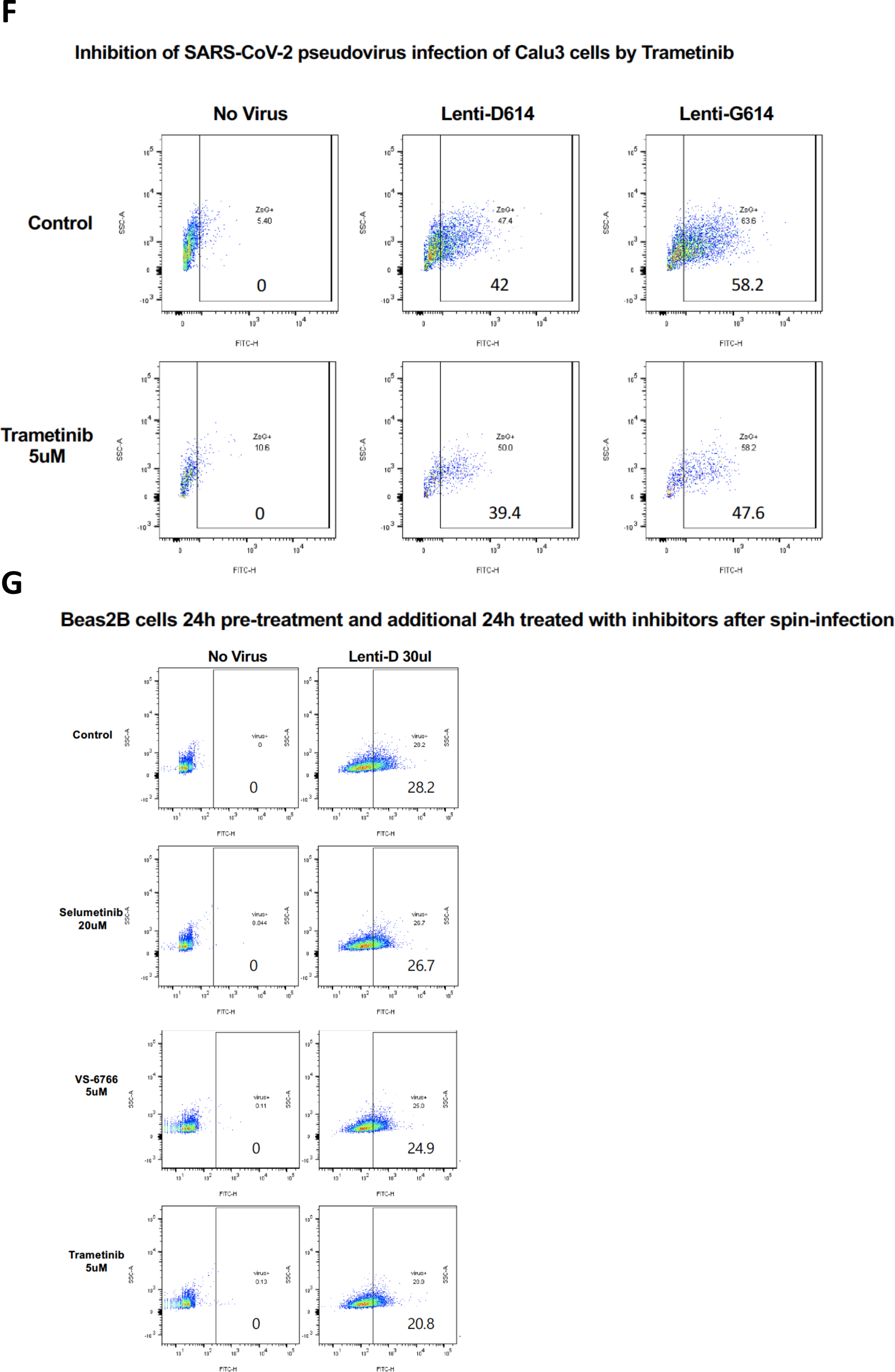
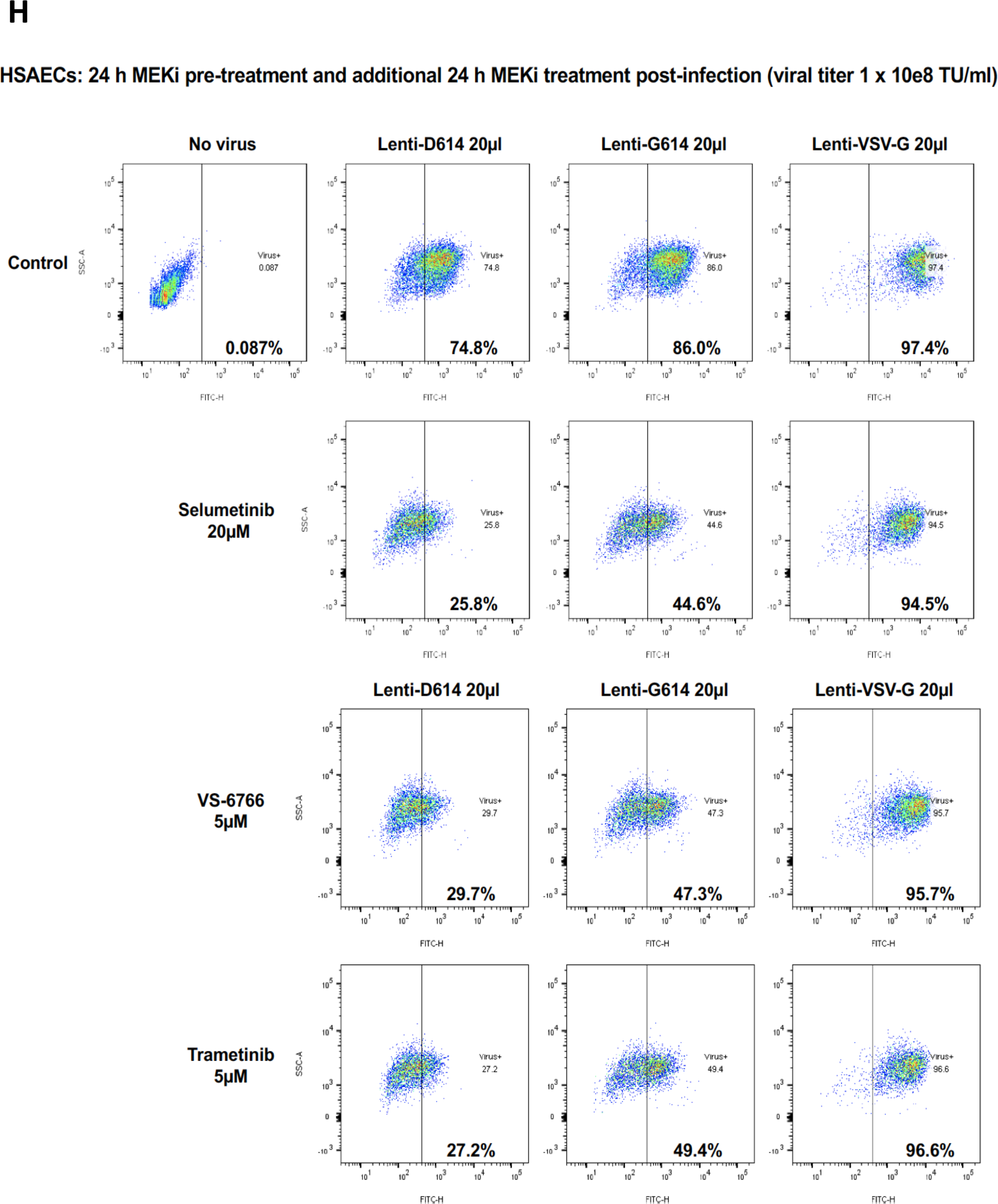
Establishment of a SARS-CoV-2 pseudovirus that expresses SPIKE protein variants on the envelope of a lentiviral core, infection of human airway epithelial cells or lung cancer cells, and demonstration of MEKi attenuation of infectivity. (A) Western blot to demonstrate FLAG-tagged S protein of SARS-CoV-2 pseudoviruses. We generated a pseudotyped SARS-CoV-2 virus which has a lentiviral core but with the SARS-CoV-2 D614 or G614 spike (S) protein on its envelope. VSV-G lentivirus was used as a negative control. Lenti-X™ p24 Rapid Titration ELISA Kit (TaKaRa) was used to determine virus titer. Equal number of lentiviral particles were analyzed on an SDS-PAGE gel followed by Western blot to detect FLAG-tagged S protein. The results suggest more G614 than D614 S protein on each viral particle. (B) Fluorescence microscopic evaluation of SARS-CoV-2 pseudovirus infection of BEAS-2B cells. SARS-CoV-2 pseudoviruses (5 x 10^6^) or VSV-G lentivirus (2 x 10^5^) were used to spin-infect BEAS-2B cells in a 6-well plate. Fluorescence microscopic images were taken 18 h after infection. (C) Flow cytometry analysis of SARS-CoV-2 pseudovirus infection of BEAS-2B cells. SARS-CoV-2 pseudoviruses (5 x 10^6^) or VSV-G lentivirus (2 x 10^5^) were used to spin-infect BEAS-2B cells in a 6-well plate. Flow cytometry analysis of ZsGreen+ cells was carried out 48 h after infection. (D) Fluorescence microscopic evaluation of SARS-CoV-2 pseudovirus infection of Calu-3 cells. SARS-CoV-2 pseudoviruses (5 x 10^6^) or VSV-G lentivirus (2 x 10^5^) were used to spin-infect Calu-3 cells in a 6-well plate. Fluorescence microscopic images were taken 18 h after infection. (E) Flow cytometry analysis of SARS-CoV-2 pseudovirus infection of Calu-3 cells. SARS-CoV-2 pseudoviruses (5 x 10^6^) or VSV-G lentivirus (2 x 10^5^) were used to spin-infect Calu-3 cells in a 6-well plate. Flow cytometry analysis of ZsGreen+ cells was carried out 48 h after infection. (F) Inhibition of SARS-CoV-2 pseudovirus infection of Calu-3 cells by Trametinib. Calu-3 cells were pre-treated with the inhibitor for 48 h, spun-infected with pseudovirus followed by another 48 h incubation with the inhibitor. Flow cytometry analysis of live cells that were ZsGreen+ was carried out. (G) Inhibition of SARS-CoV-2 pseudovirus infection of BEAS-2B cells by MEKi. BEAS-2B cells were pre-treated with MEKi (doses as indicated) for 24 h, spun-infected with pseudovirus followed by another 24 h incubation with the various MEKi. Flow cytometry analysis of live cells that were ZsGreen+ was carried out. (H) Inhibition of SARS-CoV-2 pseudovirus cell entry of Human Small Airway Epithelial Cells (HSAECs) by MEKi. HSAEC were grown on 12-well plates till 80% confluence, pre-treated with the inhibitors at the indicated concentrations for 24 hr, spun-infected with the pseudoviruses followed by another 24 hr incubation with the inhibitors. All MEKi significantly blocked pseudovirus cell entry of HSAECs while having no effect on the pantropic VSV-G lentivirus cell entry. Overall cell survival was more than 75% as determined by DAPI-in flow cytometry. The percentage of ZsGreen+ live cells was analyzed by using the FlowJo software version 10 (FlowJo, LLC, Ashland, OR).

We performed an additional experiment evaluating the effects of the three MEK inhibitors on the D and G SPIKE variants in pseudovirus infectivity of human small airway epithelial cells (HSAECs). All MEKi blocked pseudovirus cell entry of HSAECs in a very significant way, while having no effect on the pantropic VSV-G cell entry (Figure 6H). Overall cell survival was more than 75%.

## Discussion

The goal of this work was to identify candidate drugs that are available for clinical use that could be tested for their efficacy in suppressing the infectivity of SARS-CoV-2 as well as disease severity in preclinical studies. Our approach was to evaluate candidate small molecules for their effects on SARS-CoV-2 infectivity factors such as ACE2, and TMPRSS2 expression while monitoring their impact on host immune suppressors of viral infection (Natural Killer cells and TRAIL-mediated cell killing) and cytokine release that has been correlated with COVID-19 disease severity. The identification of MEK inhibitors as having multiple favorable activities in these assays could support their clinical testing to suppress SARS-CoV-2 infection and if combined with an anti-viral agent such as remdesivir there could be enhancement of anti-viral efficacy under conditions where both infectivity was suppressed and innate immune responses were enhanced.

We envisioned the potential to test these ideas in the clinic in patients with early COVID-19 infection where it may be possible to control the progression and spread of the infection throughout the body. Our hypotheses were that suppression of viral entry into the cell could be achieved through inhibition of ACE2 and TMPRSS2 expression, that blocking viral infection will reduce the spread of SARS-CoV-2 and allow the innate immune system and antivirals such as remdesivir to more effectively suppress viral infection, and that combinations of drugs that reduce ACE2 and TMPRSS2 may be helpful in addressing unexpected effects of remdesivir on ACE2 expression or hydroxychloroquine on active TMPRSS2.

We further explored the relationship between MEK and ACE2 given prior literature that SARS coronavirus SPIKE protein can increase ACE2 and stimulate the MAPK pathway and that MAPK pathway regulates ACE2. We reasoned that if MEKi impact on ACE2 expression, could they be used to attenuate early infection? Moreover, could MEKi be combined with remdesivir to improve its antiviral efficacy? We observed that several MEK inhibitors suppress ACE2 expression at nontoxic doses either alone or in the presence of remdesivir. We observed unexpectedly that under some experimental conditions, remdesivir increases ACE2 promoter activity, mRNA expression and protein expression, and this is suppressed when remdesivir is combined with MEKi such as VS-6766.

It is important to note and emphasize that there is variability and heterogeneity in the extent of ACE2 suppression by MEKi and in the increase of ACE2 in remdesivir-treated cells. While we observed correlations between pERK and ACE2 expression under different experimental conditions, we do not currently understand the variability in drug effects in different cells. The approaches included serum starvation and stimulation, addition of recombinant SPIKE protein or use of drugs on log-phase cells under their normal growth conditions. The recombinant SPIKE protein experiments lack negative controls (could be non-specific) and we do not know why both S1 and S2 stimulate pERK since S2 is not known to have a receptor-binding domain. Examples of more subtle effects include subtle increase of ACE2 protein in BEAS-2B or H522 cells after remdesivir, while in combination with VS-6766 there is reduction of both glycosylated and nonglycosylated ACE2. In some cells, the reduction of ACE2 is appreciated while an increase by remdesivir is not observed. This is a limitation of the work, as is lack of direct evidence that MEKi attenuate SARS-CoV-2 infection of human cells. Our evidence is indirect, and the effects are predicted based on current knowledge of SARS-CoV-2 infectivity factors, and consistent with recent results showing SARS-Cov-2 effects on the kinome including MAPK p38 activation [45]. Our results showing infection of SARS-CoV-2-S pseudovirus of human bronchial epithelial cells, human small airway epithelial cells, or lung cancer cells provide an experimental model system to discover or test therapeutics with potential to block coronavirus infection. The observed effect of MEKi to reduce SARS-CoV-2-S pseudovirus infection of human cells is consistent with our other evidence that MEKi may attenuate coronavirus infectivity factors to inhibit infection.

In pursuit of a therapeutic agent that could attenuate cytokine storm while reducing viral infectivity and boosting NK cells activity, we found that VS-6766 decreases G-CSF and other cytokines. These cytokines of interest were increased in COVID-19-(+) patient plasma samples in our study. The combination of remdesivir and VS-6766 was not associated with increased cytokine expression at nontoxic doses of the drugs. The MEKi plus remdesivir drug combinations do not block NK-mediated cell killing and in fact the MEKi stimulate NK killing activity towards target cells. Moreover, the drug combinations do not inhibit TRAIL-mediated killing of target cells. The observed stimulation of NK cell killing of target cells by MEK inhibitors is a novel finding that is relevant and important not only in the context of the current work focused on COVID-19 but also for cancer therapy mechanisms. Additional experiments will need to further evaluate the role of NK cells in anti-tumor efficacy *in vivo,* for example by using immune-depletion approaches [30].

Our results support the idea that MEK inhibitors as a drug class may suppress COVID-19 infectivity factors while allowing (or in some cases boosting) NK-mediated (but not T-cell mediated) killing of target cells and suppressing inflammatory cytokines. The results support the idea that MEKi could be tested in the clinic to suppress early COVID-19 infection and that in combination with an anti-viral such as remdesivir, MEKi may provide a means to lessen the infection spread which may potentiate anti-viral effects. Limitations of this work include a focus on host factors without the presence of actual SARS-CoV-2 infection, although some experiments employed recombinant SPIKE protein fragments. However, we did create a SARS-CoV-2-S pseudovirus bearing D614 or G614 SPIKE protein variants on the envelope of a lentiviral core, demonstrated infection of human airway epithelial cells or lung cancer cells, and showed that MEKi attenuate the viral infectivity. We are conducting additional experiments evaluating the effects of MEKi on pseudovirus infection of other cell lines and also investigating the impact of MEKi on NK cell killing of cells already infected by the pseudovirus.

Our understanding of the pathogenesis of COVID-19 has evolved rapidly over the course of the pandemic [24, 46, 47]. Several classes of drugs are being evaluated for the management of COVID-19: antivirals, antibodies, anti-inflammatory agents, targeted immunomodulatory therapies, anticoagulants, and antifibrotics. However, no proven drug for the treatment of COVID-19 is currently available [48]. Our inhibitors capable of both inhibiting viral infection and modulating immune responses may synergize to block the disease at multiple stages over its natural history. Our approach with lentiviral vector based pseudovirus has its limitations. For example, how the display of S proteins on a heterologous virus impacts viral entry compared to infectious SARS-CoV-2 is not known. Also, the ability of lentivirus to elicit immune response in infected cells is limited. Further studies of SARS-CoV-2 virus cell entry and its inhibition with a replication competent surrogate coronavirus [49, 50] are warranted.

An additional prior literature supports the potential use of MEKi to reduce systemic inflammation and enhance the immune response in vivo. For example, in the setting of sepsis, MEKi prevents endotoxin-induced kidney injury in mice [51]. Studies have shown that MEKi trametinib in a cecal puncture-sepsis model reduced cytokines as well as kidney, liver, and muscle injury *in vivo* [52]. In chronic obstructive pulmonary disease (COPD), MEK1/2 inhibition has an anti-inflammatory effect in human alveolar macrophages while promoting increased bacterial killing in neutrophils [53]. MEKi selumetinib has been previously observed to reduce IL-6 levels in a Lewis lung carcinoma model although it did not protect against cachexia [54]. MEKi have also been shown to not inhibit dendritic cell priming by T-cells and to promote synergistic anti-tumor immunity when combined with an immunostimulatory CD40 agonist [55]. These findings are consistent with our observations and add further *in vivo* evidence regarding the anti-inflammatory and immune-boosting effects of MEKi that we suggest are relevant to pursue in suppression of early COVID-19 disease.

Based on the data in this manuscript it may be reasonable to consider further preclinical experiments as well as clinical translation of the MEKi results. Some of the open questions include a more detailed understanding of how the MAPK pathway activates ACE2, more direct evidence for effects of MEKi on actual SARS-CoV-2 infectivity of human cells, and more evidence for their effects on COVID-19 infection spread in preclinical models. In the clinic, it may be reasonable to test MEKi such as VS-6766 or trametinib in COVID-19 infected but less severely ill patients to test the idea that MEKi could keep the infection from getting worse while allowing the body’s NK cells and innate immune mechanisms to more effectively attack virally infected cells prior to severe infection. Consideration could be given to evaluation of MEKi −/+ antiviral agents such as remdesivir given results suggesting potentially favorable drug interactions that may allow suppression of infectivity, suppression of inflammatory cytokines, stimulation of NK cell (but not T-cell) activity, and lack of suppression of TRAIL-mediated cytotoxicity. These effects may help antiviral agents achieve more potent disease suppression to attenuate or block COVID-19 infection that may be of use as a therapeutic approach in patients with early or less severe COVID-19 disease.

## Materials and Methods

### Human Plasma Samples

COVID-19 (+) human plasma samples were received from the Lifespan Brown COVID-19 biobank at Rhode Island Hospital (Providence, Rhode Island). All patient samples were deidentified but with available clinical information as described in the manuscript. The IRB study protocol “Pilot Study Evaluating Cytokine Profiles in COVID-19 Patient Samples” did not meet the definition of human subjects research by either the Brown University or the Rhode Island Hospital IRBs. Normal, healthy, COVID-19 (−) samples were commercially available form Lee BioSolutions (991-58-PS-1, Lee BioSolutions, Maryland Heights, Missouri). All samples were thawed and centrifuged to remove cellular debris immediately before the assay was ran.

### Cytokine Measurements of Culture Supernatants and Plasma Samples

A MilliPlex MILLIPLEX® MAP Human Cytokine/Chemokine/Growth Factor Panel A-Immunology Multiplex Assay (HCYTA-60K-13, Millipore Sigma, Burlington, Massachusetts) was run on a Luminex 200 Instrument (LX200-XPON-RUO, Luminex Corporation, Austin, Texas) according to the manufacturer’s instructions. Production of granulocyte colony-stimulating factor (G-CSF), interferon gamma (IFNγ), interleukin 1 alpha (IL-1α), interleukin-1 receptor antagonist (IL-1RA), IL-2, IL-6, IL-7, IL-12, interferon γ-induced protein 10 (IP-10), monocyte chemoattractant protein-1 (MCP-1), macrophage colony-stimulating factor (M-CSF), macrophage inflammatory protein-1 alpha (MIP-1α), and tumor necrosis factor alpha (TNFα) in the culture supernatant were measured. All samples were run in triplicate.

### Cell lines and culture conditions

Normal human primary small airway epithelial cells HSAEC, normal human bronchial epithelial cells BEAS-2B, normal human lung fibroblast MRC-5, human NSCLC cells H1975, H1299, Calu-3, Calu-6, human mesothelioma cells MSTO-211H, NSCLC patient-derived cell line, human natural killer cells NK-92, normal human colon epithelial cells CCD 841 CoN and human colorectal cancer cells HT-29, HCT116 were used in this study. HSAEC was cultured in Airway Epithelial Cell Basal Medium (ATCC® PCS-300-030™) supplemented with Bronchial Epithelial Cell Growth Kit (ATCC® PCS-300-040™). BEAS-2B was cultured in BEGM™ Bronchial Epithelial Cell Growth Medium BulletKit™ (Lonza Catalog No. CC-3170). TALL-104 T cells were from the ATCC (ATCC® CRL-11386™) and were used as we recently described [31]. MRC-5, Calu-3, Calu-6 and CCD 841 CoN cells were cultured in Eagle’s Minimum Essential Medium supplemented with 10% fetal bovine serum (FBS). H1975, H1299, MSTO-211H and NSCLC patient-derived cell line were cultured in RPMI-1640 medium supplemented 10% FBS. NK-92 cells were cultured in Alpha Minimum Essential medium without ribonucleosides and deoxyribonucleosides but with 2 mM L-glutamine and 1.5 g/L sodium bicarbonate supplemented with 0.2 mM inositol; 0.1 mM 2-mercaptoethanol; 0.02 mM folic acid; 100 U/ml recombinant IL-2, 12.5% horse serum and 12.5% FBS. HCT116 and HT-29 were cultured in McCoy’s 5A (modified) medium supplemented 10% FBS. All cell lines were incubated at 37 °C in humidified atmosphere containing 5% CO2. Media containing 1% serum was used for serum starvation experiments. Cell lines were authenticated and tested to ensure the cultures were free of mycoplasma infection.

### Collection of Culture Supernatants Used in Cytokine Measurements

Cells were plated at 3.5 × 10^4^ cells in a 48 well plate (Thermo Fisher Scientific, Waltham, MA) in CM and incubated at 37°C with 5% CO2. At 24 h after plating, almost all the tumor cells were adherent to the bottom of the flask and then the CM was completely replaced. Subsequently, the culture supernatants were collected after another 48 hr of incubation, centrifuged to remove cellular debris, and then frozen at −80°C until the measurement of cytokines was performed.

### Transfections and reporter assays

Human ACE2-Luc promoter constructs containing 1119, 252, and 202 base pairs of the ACE2 promoter linked to firefly luciferase were obtained from Addgene. Human tumor cell lines were transfected with each ACE2-promoter luciferase-reporter using lipofectamine 2000 as described in the protocol (Invitrogen, USA). At 24 hours after the transfection, the cells were trypsinized and seeded into 96-well black plates at a density of 3 × 10^4^ cells/well. The next day, cells on the 96-well plate were treated with drugs for 24 hours. The luciferase activity was evaluated by bioluminescence imaging in cells using the IVIS imaging system (Xenogen, Alameda, CA). DMSO treatment was used as a negative control in each screened plate. The bioluminescence in each treatment was normalized to DMSO treatment.

### Cell viability assays

Cells were plated at a density of 3 x 10^3^ cells per well of a 96-well plate. Cell viability was assessed using the CellTiter Glo assay (Promega). Cells were mixed with 25 μl of CellTiter-Glo reagents in 100 μl of culture volume, and bioluminescence imaging was measured using the Xenogen IVIS imager.

### Western blots and antibodies

Immunoblotting for proteins was performed using the following antibodies: Cell Signaling Technology ACE2 Antibody #4355, Abnova ACE2 polyclonal antibody #PAB13444, Santa Cruz Biotechnology ACE2 Antibody (E-11) #sc-390851, Sigma Anti-TMPRSS2 Antibody, clone P5H9-A3 #MABF2158, Sigma Anti-IL6 antibody produced in rabbit #SAB1408591, Santa Cruz Biotechnology TMPRSS2 Antibody (H-4) #sc-515727, TMPRSS2 (EMD Millipore #MABF2158), Cell Signaling Technology Phospho-p44/42 MAPK (Erk1/2) (Thr202/Tyr204) (D13.14.4E) XP® Rabbit mAb #4370, Cell Signaling Technology p44/42 MAPK (Erk1/2) Antibody #9102, Ran (BD Biosciences #610341), Caspase-8 (Cell Signaling #9746), Sigma Monoclonal Anti-β-Actin antibody produced in mouse #A5441. Cell Signaling Rb (4H1) Mouse mAb #9309, and Phospho-Rb (Ser807/811) (D20B12) XP® Rabbit mAb #8516. Primary antibodies are diluted according to their datasheets. Proteins were extracted from cells in denaturing sample buffer and an equal amount of protein lysate was electrophoresed through 4-12% SDS-PAGE (Invitrogen) then transferred to PVDF membranes. The PVDF membrane was blocked with 5% non-fat milk (Sigma) in 1x PBS. The primary antibodies indicated in the figures were incubated with the transferred PVDF in blocking buffer at 4°C overnight. Antibody binding was detected on PVDF with appropriate Pierce HRP-conjugated secondary antibodies by the Syngene imaging system. Invitrogen Goat anti-Rabbit IgG (H+L) Secondary Antibody, HRP # 31460 and Goat anti-Mouse IgG (H+L) Secondary Antibody, HRP # 31430 were diluted 1:5000 in 2.5% non-fat milk.

### qRT-PCR methods and primers

Total RNA was isolated from cells using the RNeasy Mini Kit (Qiagen). Reverse transcription was performed with random primers using the SuperScript II First-Strand Synthesis System (Invitrogen). qRT-PCR reactions used SYBR Green Master Mix with the Real-Time PCR Detection systems (Bio-Rad). Primers for ACE2 were obtained from Origene.

Forward Sequence: 5’-TCCATTGGTCTTCTGTCACCCG-3’.

Reverse Sequence: 5’-AGACCATCCACCTCCACTTCTC-3’.

The ACE2 mRNA gene expression levels were normalized with GAPDH.

### NK-cell co-culture system and microscopic imaging for data analysis

Tumor cells were plated and allowed to grow for 48 hours in their culture media before addition of NK cells. Green fluorescent tumor cells were co-cultured with NK-92 cells [30] at a 1:1 effector target cell ratio (E:T) imaged at various timepoints. Cells were treated with indicated drugs for several hours (drugs added at time of NK cell addition) as indicated in the figures. A total of 1 μM red fluorescent ethidium homodimer (EthD-1) was added at the beginning of drug and NK cell incubation to detect dead cells (Invitrogen, Waltham, MA). For the quantification of dead/live cells, fluorescence microscopy was used to take images at 10x magnification. The number of red/green color cells in random fields was determined using thresholding and particle analysis in the Fiji modification of ImageJ and expressed as a dead/live cell ratio. At least 100 cells were evaluated per sample, with 3 independent replicates.

### TRAIL cell killing assays

Cells were plated at a density of 5 x 10^5^ per well of a 6-well plate. After 16 hours of incubation at 37 degrees Celsius in 5% CO2, cells were treated with 5 μM remdesivir (RDV), VS-6766, or the combination. After 24 hours, cells were treated with TRAIL (50 ng/mL) for an additional 4 hr. Western blots evaluating cleaved caspase 8 were performed using Cell Signaling antibody (#9746). Ran was probed with BD Biosciences antibody (#610341) as a loading control.

### SARS-CoV-2 pseudovirus production, quantification, and infection

We used a lentiviral packaging system to produce a replication incompetent SARS-CoV-2 pseudovirus. Briefly, HEK293T cells at 75% confluency were co-transfected with the backbone vector pHAGE-fullEF1α-ZsGreen-IRES-Puro(R), plasmids expressing lentiviral proteins Tat, Rev and Gag/Pol, and plasmids expressing D614 or G614 S protein (a gift from Dr. Hyeryun Choe, The Scripps Research Institute, Jupiter, FL). A plasmid expressing VSV-G protein instead of the S protein was used to generate a pantropic control lentivirus. The SARS-CoV-2 S protein gene used in the production of pseudoviruses was codon-optimized and synthesized by Integrated DNA Technologies based on the protein sequence (GenBank YP_009724390). The S protein gene is fused to the FLAG tag at its C-terminus. Pseudovirus containing culture supernatants were collected at 48 hr and 72 hr post transfection, cleared through 0.45 μm filters, and concentrated using ultra-centrifugation, aliquoted and frozen at −80°C immediately.

Lenti-X^™^ p24 Rapid Titration ELISA Kit (TaKaRa) was used to determine virus titer. Equal number of lentiviral particles were analyzed on an SDS-PAGE gel followed by Western blot to detect FLAG-tagged S protein. SARS-CoV-2 pseudoviruses (5 x 10^6^) or VSV-G lentivirus (2 x 10^5^) were used to spin-infect (931 g for 2 hr at 30°C) Calu-3 or BEAS-2B cells in a 6-well plate. Fluorescence microscopic images were taken 18 hr after infection. Flow cytometry analysis of ZsGreen+ cells was carried out 48 hr after infection on a BD LSRII flow cytometer and with the FlowJo software. To test the inhibitors, Calu-3 or BEAS-2B cells were pre-treated with the inhibitors for 48 hr, spun-infected with pseudovirus followed by another 48 hr incubation with the inhibitors. Flow cytometry analysis of live cells that were ZsGreen+ was carried out.

### Statistical methods

The statistical significance of differences between groups was determined using the Student t test. The minimal level of significance was P < 0.05. Following symbols * and ** represent, P < 0.05 and P < 0.01, respectively.

## Acknowledgements

The work was supported by a Brown University COVID-19 Seed Grant (to W.S.E-D.), and the Mencoff Family Professorship at Brown University (W.S.E-D.). W.S.E-D. is an American Cancer Society Research Professor. O.L. was supported in part by a grant from the National Institutes of Health (P20 GM119943). The COVID-19 Biobank through which plasma samples were obtained was supported by Institutional Development Award Number U54GM115677 from the National Institute of General Medical Sciences of the National Institutes of Health, which funds Advance Clinical and Translational Research (Advance-CTR). The content is solely the responsibility of the authors and does not necessarily represent the official views of the National Institutes of Health.

## Supplementary Figure Legends

**Figure S1.**
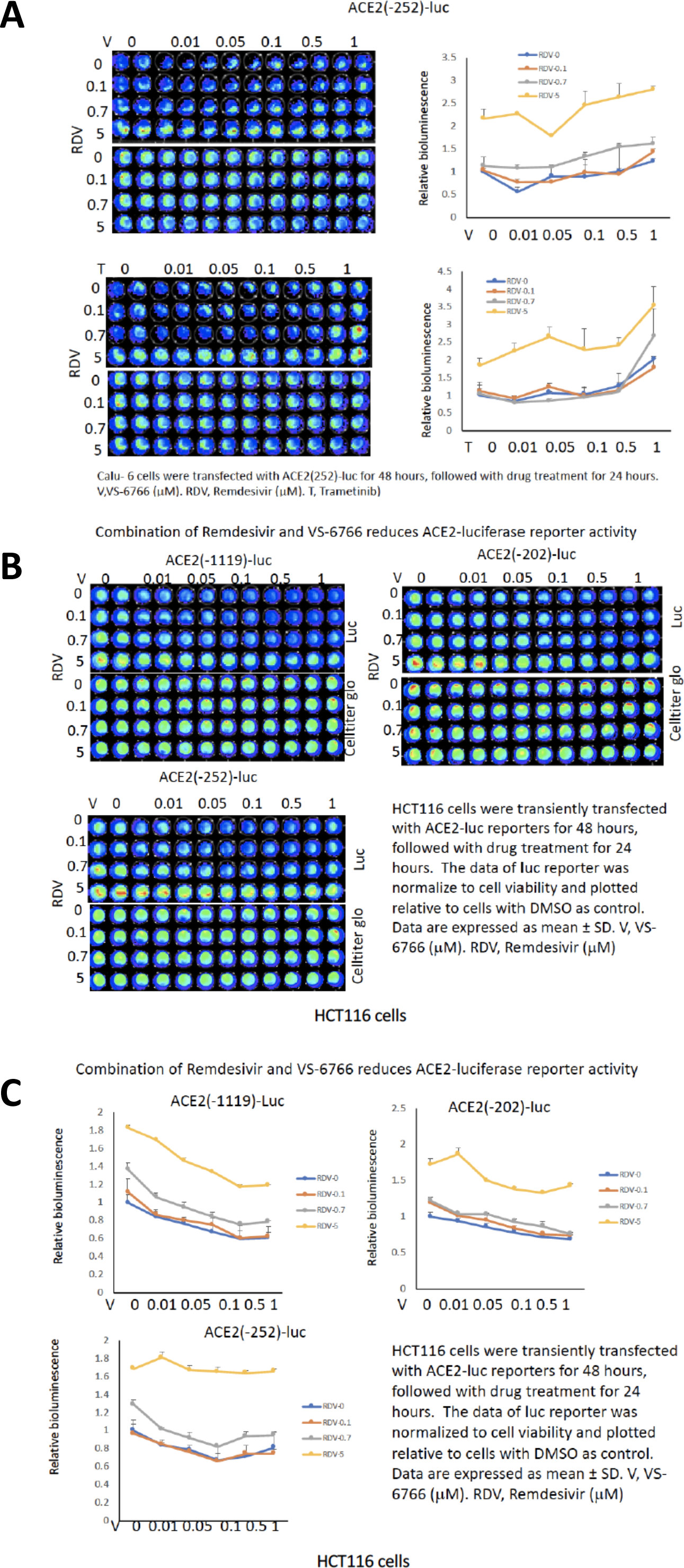
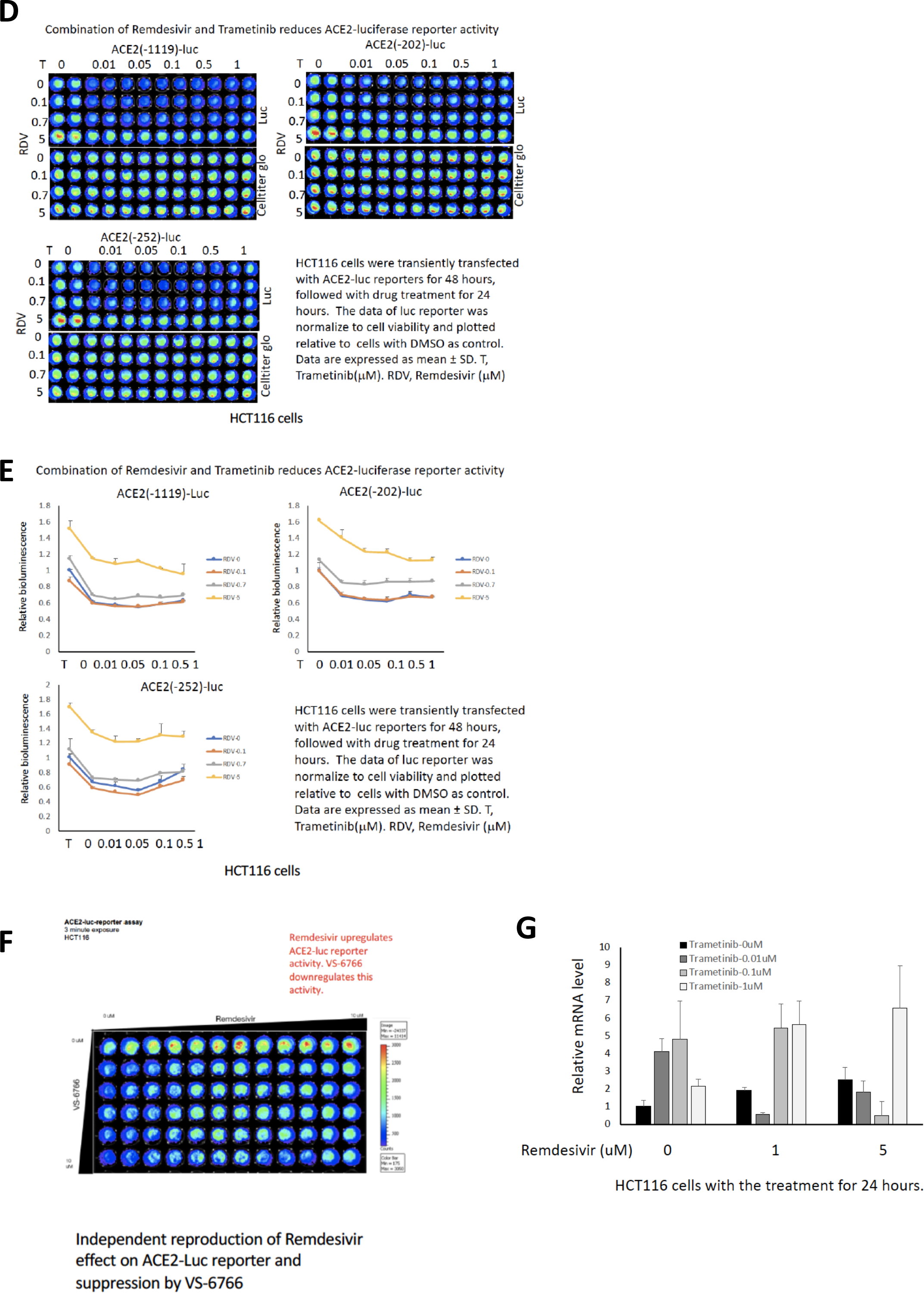
Modulation of multiple different ACE2-promoter luciferase reporters and ACE2 mRNA expression by remdesivir and MEKi in different cell lines. (A) ACE2(−252) luc reporter assay in Calu-6 cells treated with remdesivir and VS-6766 (upper panel), or remedisivir and trametinib (lower panel). (B) ACE2-luc reporter assay in HCT116 cells treated with remdesivir and VS-6766 for 24 hours. (C) The relative bioluminescence value in (B). (D) ACE2-luc reporter assay in HCT116 cells treated with remdesivir and trametinib for 24 hours. (E) The relative bioluminescence value in (D). (F) ACE2(−1119)-Luc reporter assay in HCT116 cells treated with remdesivir and VS6766. The data from the luciferase reporter assays (A-F) was normalized to cell viability and plotted relative to cells with treated with DMSO as a control. Data are expressed as mean ± SD. V, VS-6766 (μM). RDV, Remdesivir (μM). T, Trametinib (μM). (G) ACE2 mRNA level in HCT116 cells treated with remdesivir and VS-6766 for 24 hours. mRNA levels were quantified by qRT-PCR. Data were normalized to GAPDH expression and plotted relative to cells treated with DMSO as a control. Data are expressed as mean ± SD.

**Figure S2.**
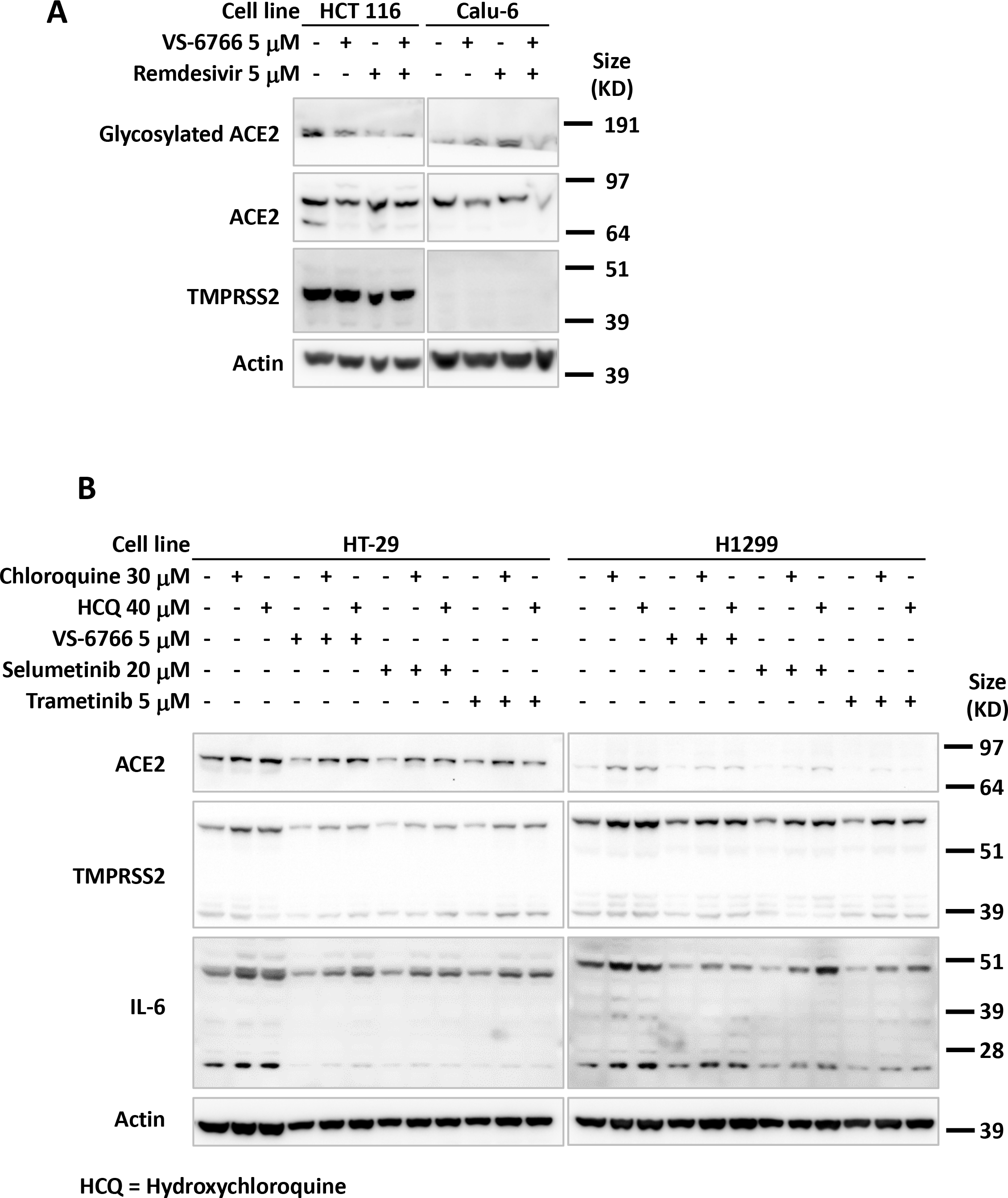
Effect of remdesivir and VS-6766 on ACE2 and TMPRSS2 in human colorectal and NSCLC cells. (A) HCT116 human CRC cells and Calu-6 human NSCLC cells were treated with remdesivir and VS-6766 at the indicated doses for 48 hours. Glycosylated ACE2, ACE2 and TMPRSS2 (full-length and Serine Protease-domain) were probed with cell signaling 4355, Abnova PAB13444, and Sigma MABF2158 antibodies. ß-Actin was probed with Sigma A5441 antibody as a loading control. (B) H1299 human NSCLC cells were treated with chloroquine, hdroxychloroquine and MEK inhibitors VS-6766, selumetinib, and trametinib at the indicated doses for 48 hours. Glycosylated ACE2, ACE2 and TMPRSS2 (full-length and Serine Proteasedomain) were probed with cell signaling 4355, Abnova PAB13444, and Sigma MABF2158 antibodies. ß-Actin was probed with Sigma A5441 antibody nas a loading control.

**Figure S3.**
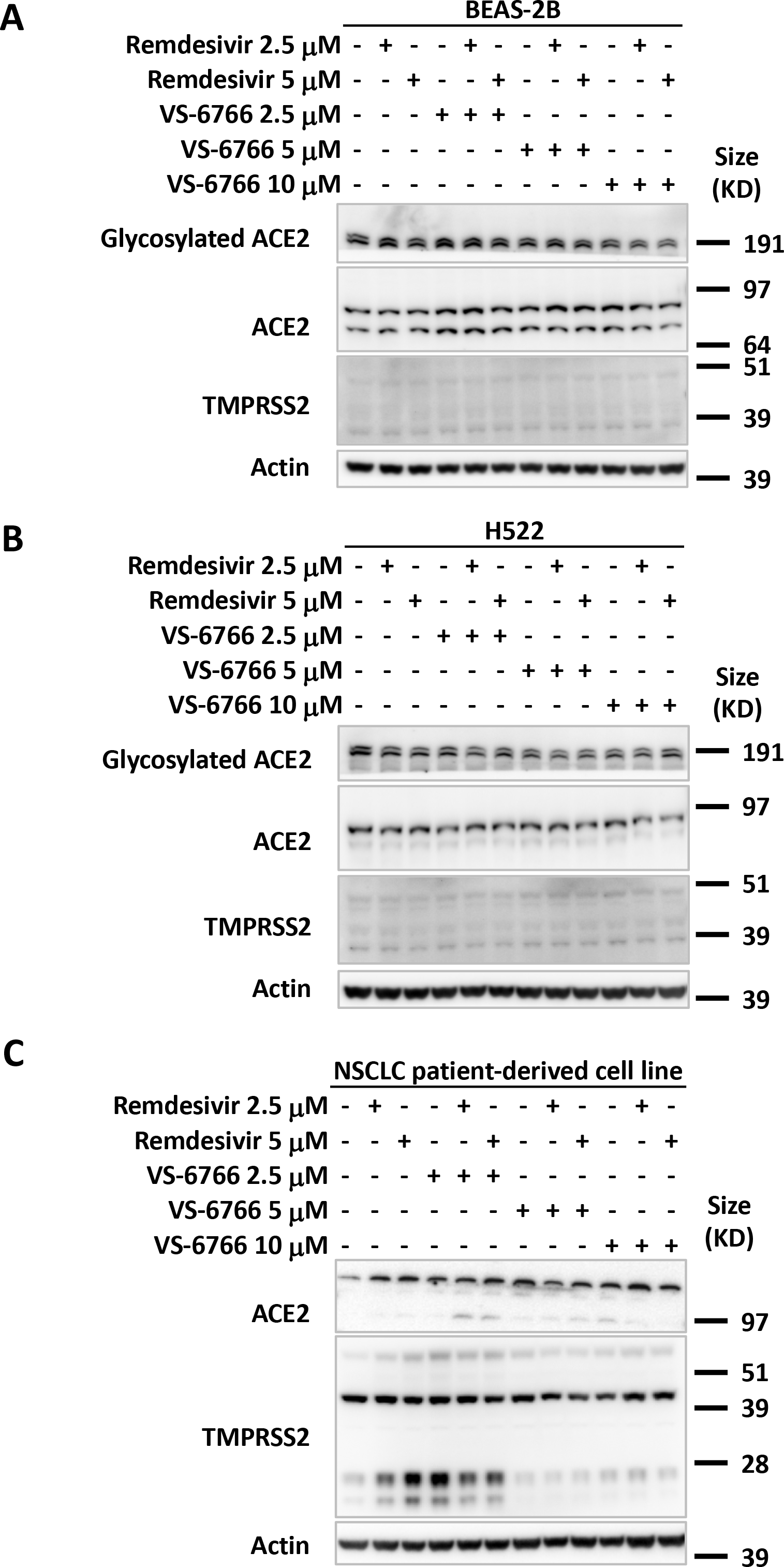
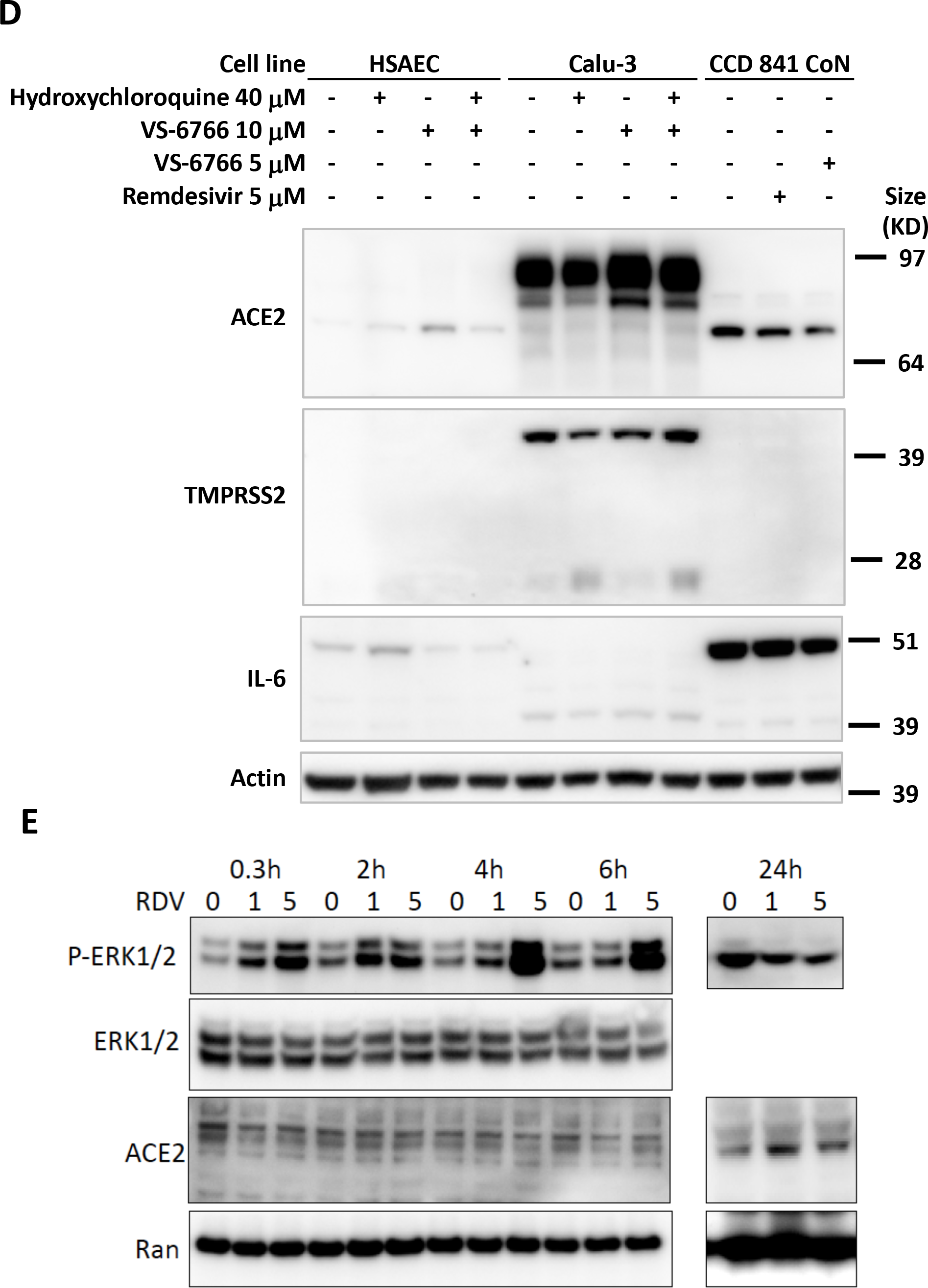
Effects of remdesivir, VS-6766 and Hydroxychloroquine on ACE2, TMPRSS2 and IL-6 in human lung and colon cells and expression of ACE2 and TMPRSS2 in cells of GI tract origin. (A) BEAS-2B normal human bronchial airway epithelial cells were treated with remdesivir and VS-6766 at the indicated doses for 48 hours. Glycosylated ACE2, ACE2 and TMPRSS2 (full-length and Serine Protease-domain) were probed with cell signaling 4355, Abnova PAB13444, and Sigma MABF2158 antibodies. ß-Actin was probed with Sigma A5441 antibody as a loading control. (B) H522 human NSCLC cells were treated with remdesivir and VS-6766 at the indicated doses for 48 hours. Glycosylated ACE2, ACE2 and TMPRSS2 (fulllength and Serine Protease-domain) were probed with cell signaling 4355, Abnova PAB13444, and Sigma MABF2158 antibodies. ß-Actin was probed with Sigma A5441 antibody as a loading control. (C) NSCLC patient-derived cell line was treated with remdesivir and VS-6766 at the indicated doses for 24 hours. ACE2 and TMPRSS2 (full-length and Serine Protease-domain) were probed with Santa Cruz sc-390851 and Sigma MABF2158 antibodies. ß-Actin was probed with Sigma A5441 antibody as a loading control. (D) Human Primary Small Airway Normal Epithelial Cells (HSAEC) and Calu-3 human NSCLC cells (Both are type II alveolar cells) were treated with hydroxychloroquine and VS-6766 at the indicated doses for 24 hours. CCD 841 CoN human colon normal epithelial cells were treated with remdesivir and VS-6766 at the indicated doses for 24 hours. Glycosylated ACE2, ACE2, TMPRSS2 (full-length and Serine Proteasedomain), IL-6 were probed with cell signaling 4355, Abnova PAB13444, Sigma MABF2158, and Sigma SAB1408591 antibodies. ß-Actin was probed with Sigma A5441 antibody as a loading control. (E) Expression of ACE2 and TMPRSS2 in cells of GI tract origin.

**Figure S4.**
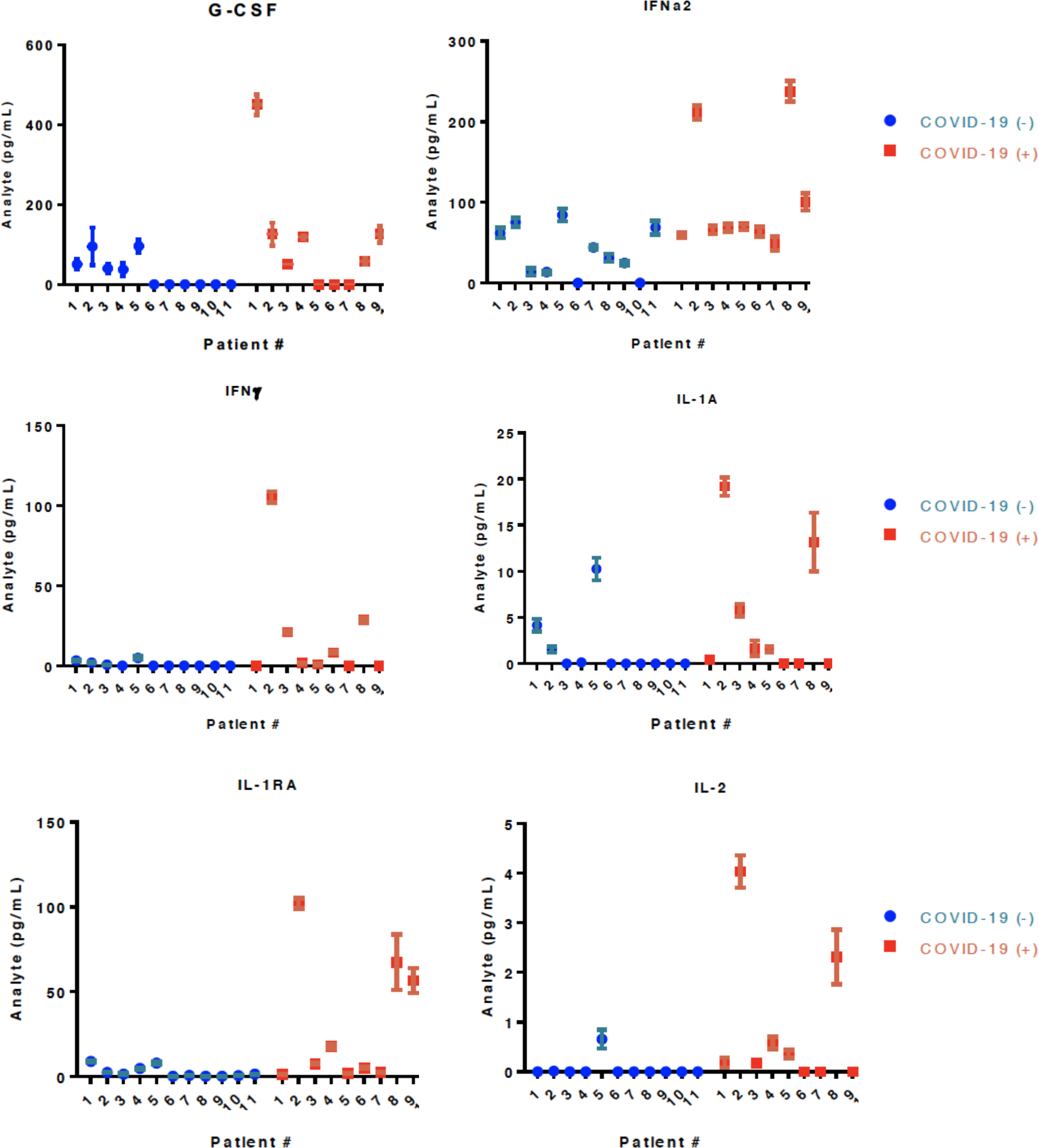
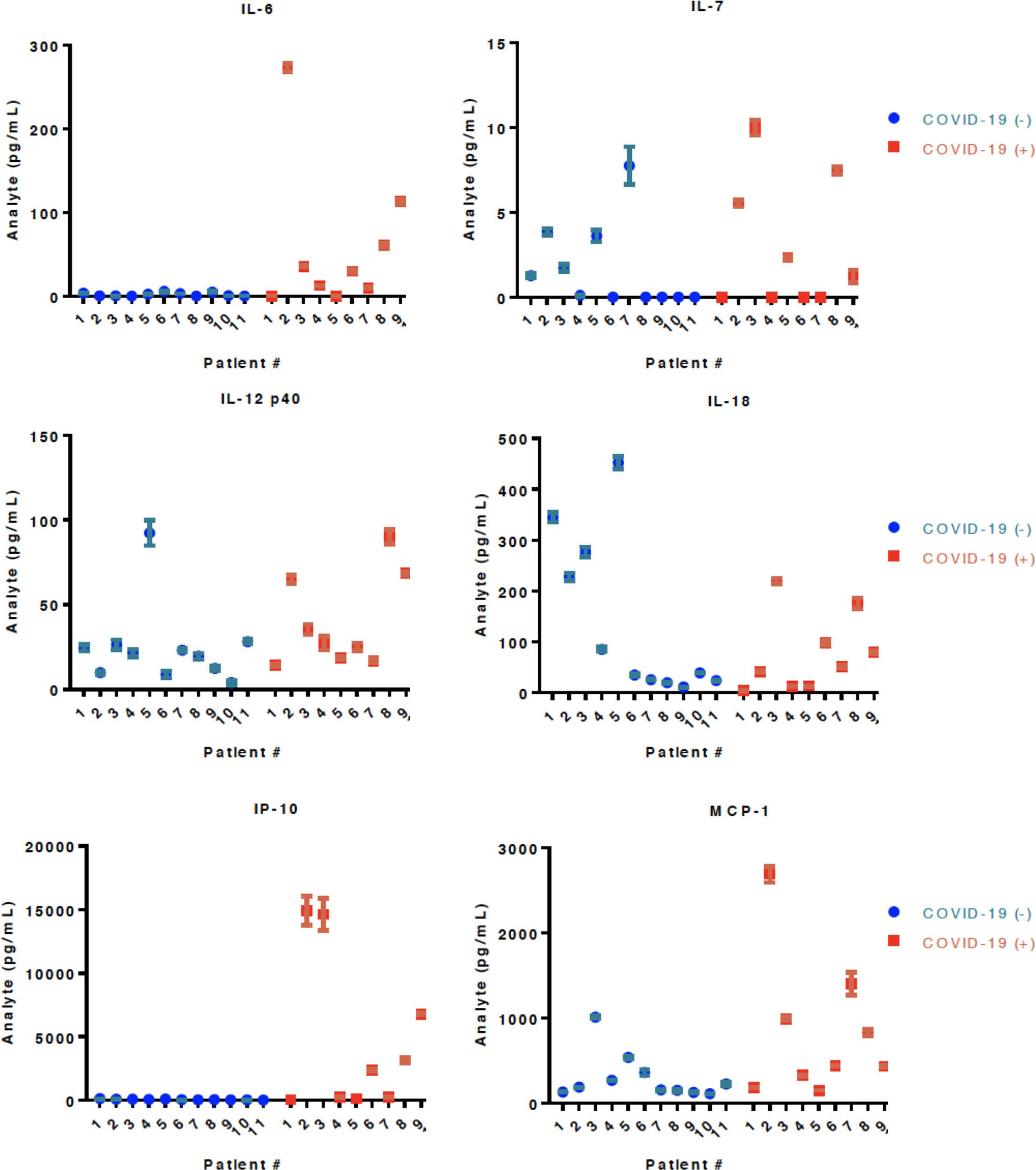
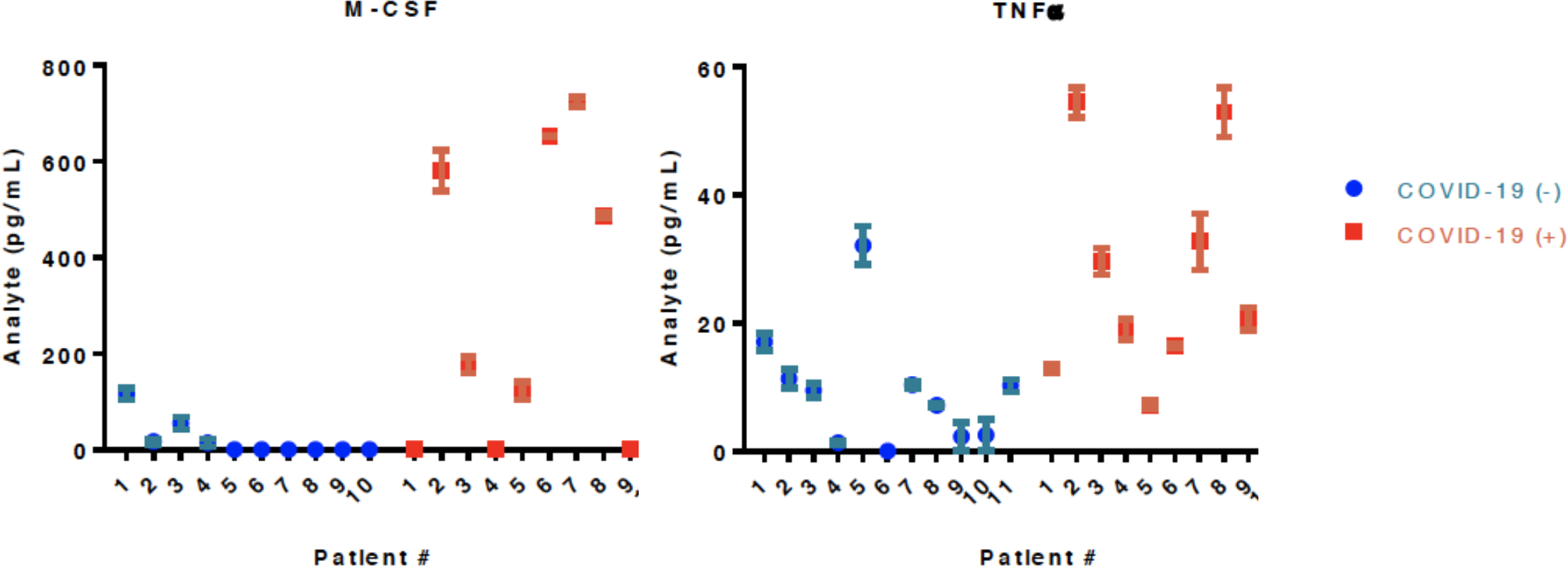
Pilot study of cytokine array profiles in COVID-19-(+) patient plasma versus control patient plasma. Cytokine levels detected in plasma are shown for individual normal (N=11) or COVID-19-(+) (N=9) patients. For the COVID-19-(+) patients the patient numbers (1-9) correspond sequentially with the numbers listed in Tables 1-5. For the COVID-19-(+) patients the cytokine levels are shown for each patient in Table 5.

**Figure S5.**
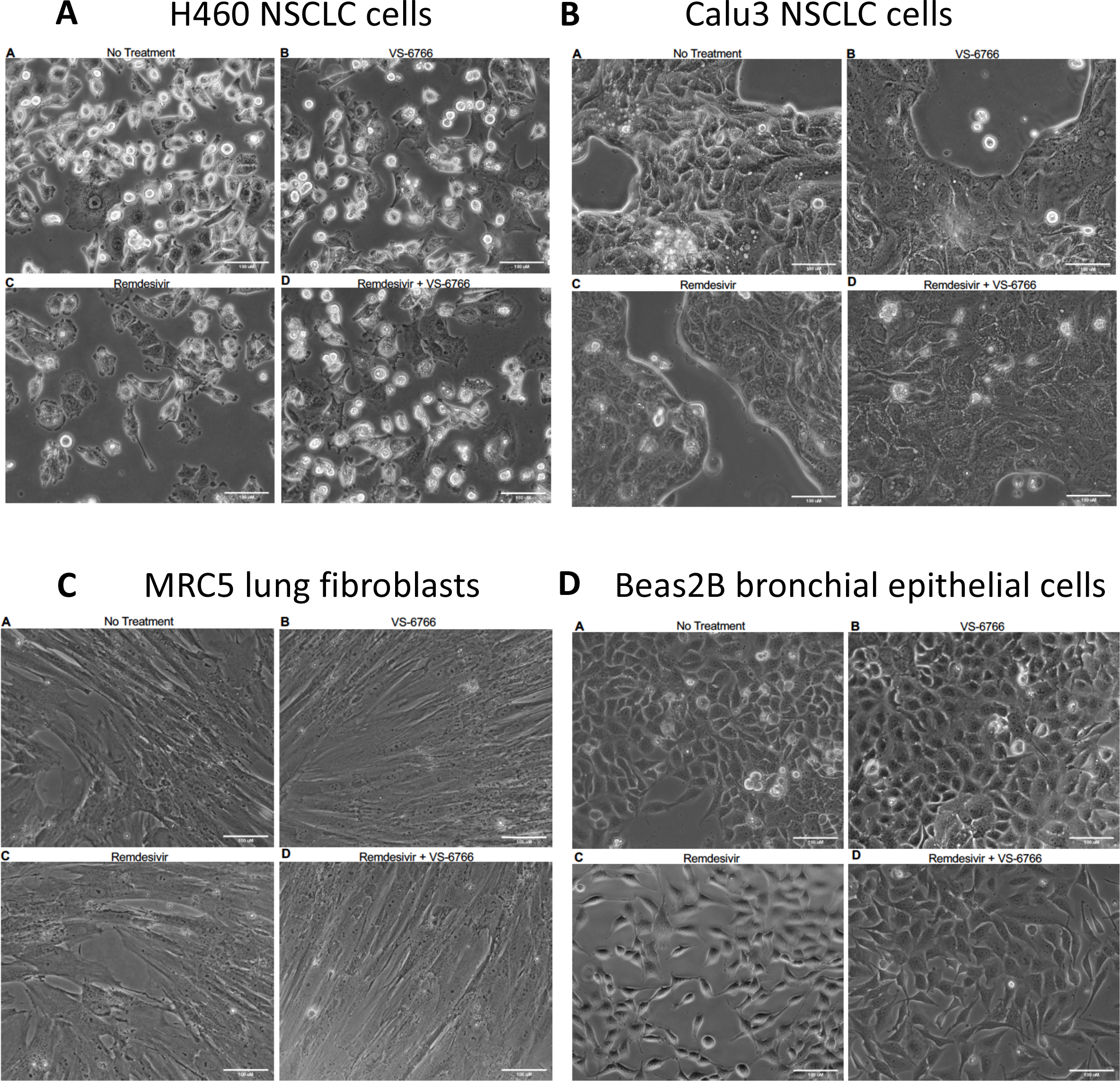
No effect observed on T cell activity by MEK inhibitor treatment at IC-50 doses. Green fluorescent SW480 tumor cells were co-cultured with TALL-104 T cells at a 1:1 effector target cell ratio (E:T) for indicated timepoints and imaged. Cells were treated with indicated drug at IC-50 doses. (A) Quantification of dead/live ratio after 4, 8 and 12 hours of treatment by drugs as indicated. P values are displayed on graph and were calculated using unpaired t tests. (B) Fluorescent microscopy of GFP+ SW480 tumor cells before and after indicated treatment conditions. Ethidium homodimer was used to visualize dead cells. 10 magnification. Scale bars indicate 100 μM. (C) Images showing GFP+ tumor target cell cytotoxic effects of MEK inhibitors alone or in addition to TALL-104 cells. Ethidium homodimer was used to visualize dead cells. (D) Quantification of tumor target cell cytotoxic effects of MEK inhibitors alone or in addition to TALL-104 cells. P values are displayed on graph and were calculated using unpaired t tests.

**Figure S6.**
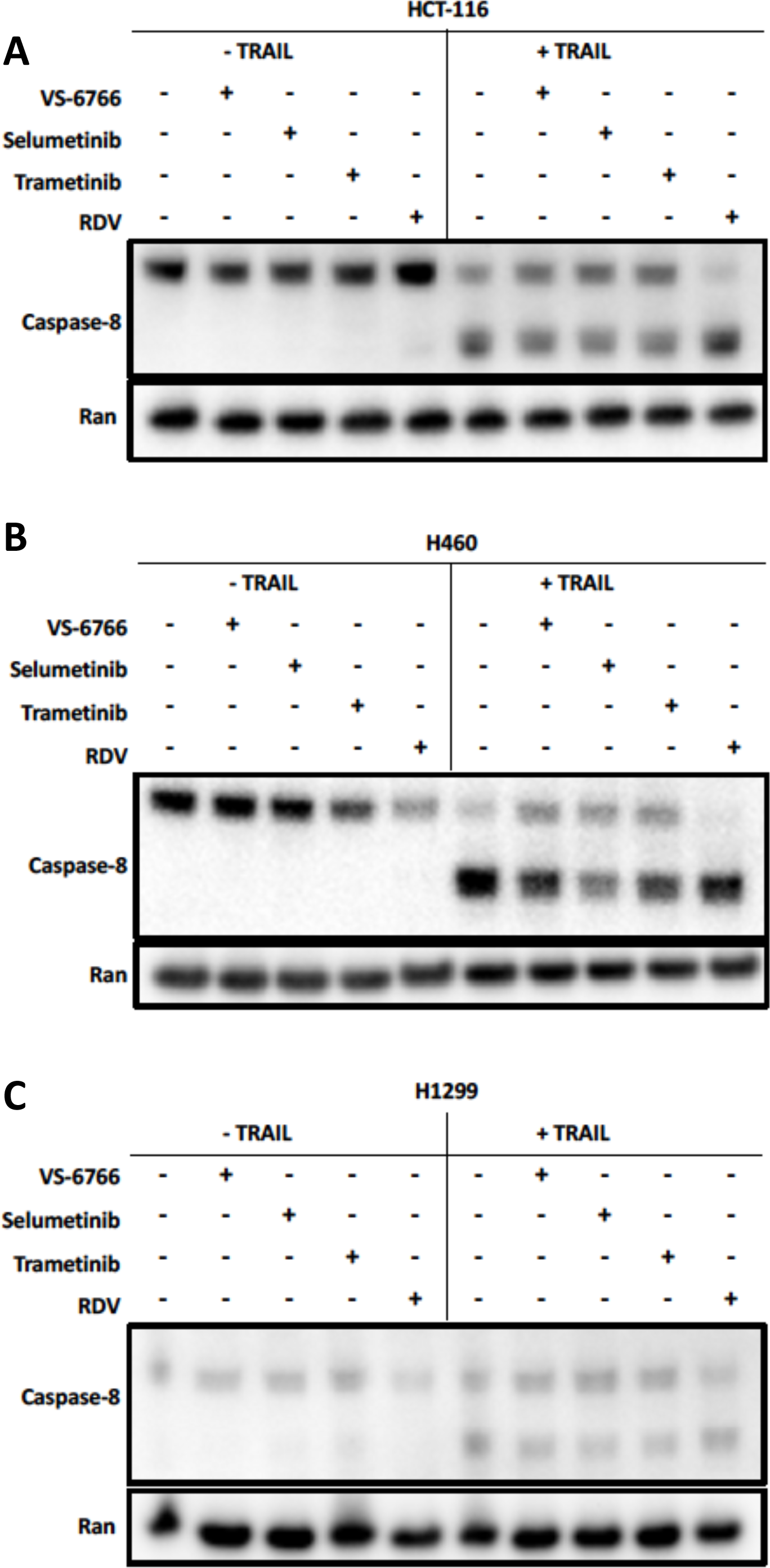
MEK inhibitors or remdesivir do not inhibit TRAIL-mediated apoptosis. Effects of VS-6766 (5 μM), selumetinib (10 μM), trametinib (5 μM), or remdesivir (5 μM) treatment for 24 hours alone or in combination with TRAIL (50 ng/mL) for 4 additional hours on cleaved caspase 8 in HCT116 colorectal cancer (A), H460 (B) or H1299 (C) lung cancer cells.

**Figure S7.**
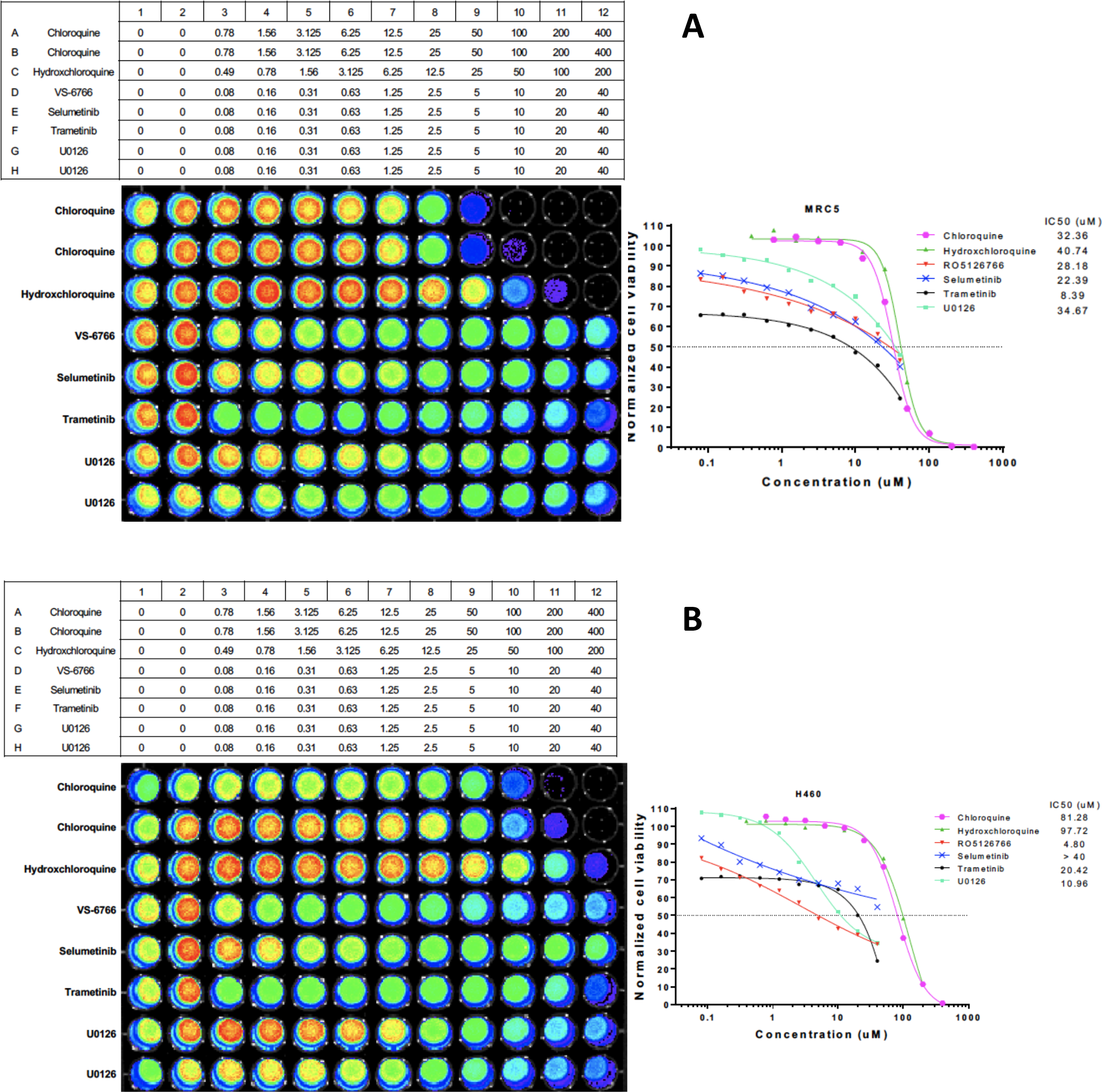
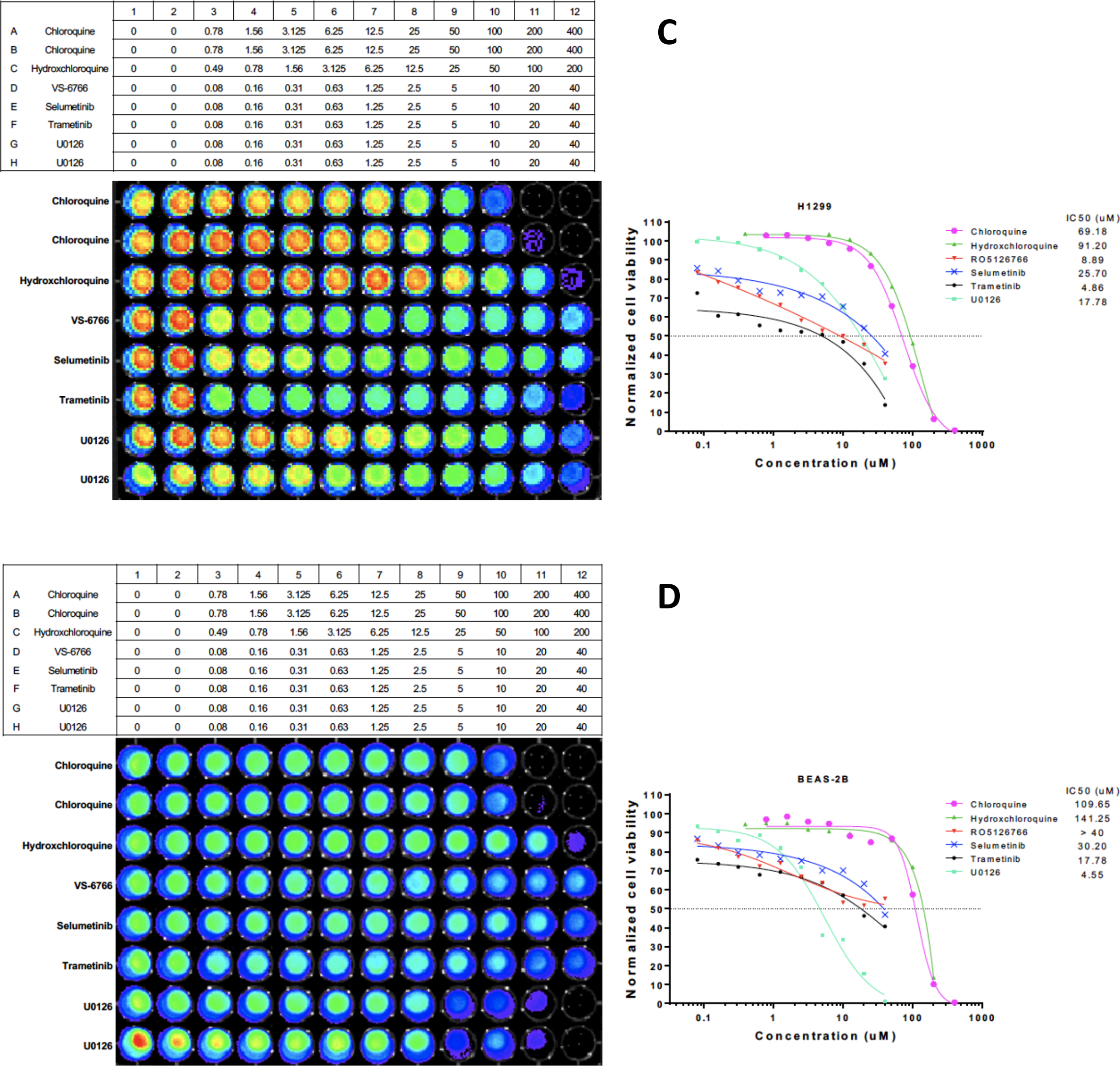
No morphological changes were observed following drug treatment with VS-6766, remdesivir or the combination in (A) H460 or (B) Calu-3 lung cancer cells, (C) normal lung fibroblasts (MRC-5) or (D) bronchial airway epithelial cells (BEAS-2B). Cells were treated for 48 hours with the following drugs in the sub-panels: (A) control; (B) 10 μM VS-6766; (C) 10 μM remdesivir; and (D) a combination treatment of 10 μM Remdesivir and 10 μM VS-6766. Micrographs were taken after 48 hours of treatment. Scale bar represents 100 μm.

**Figure S8.**
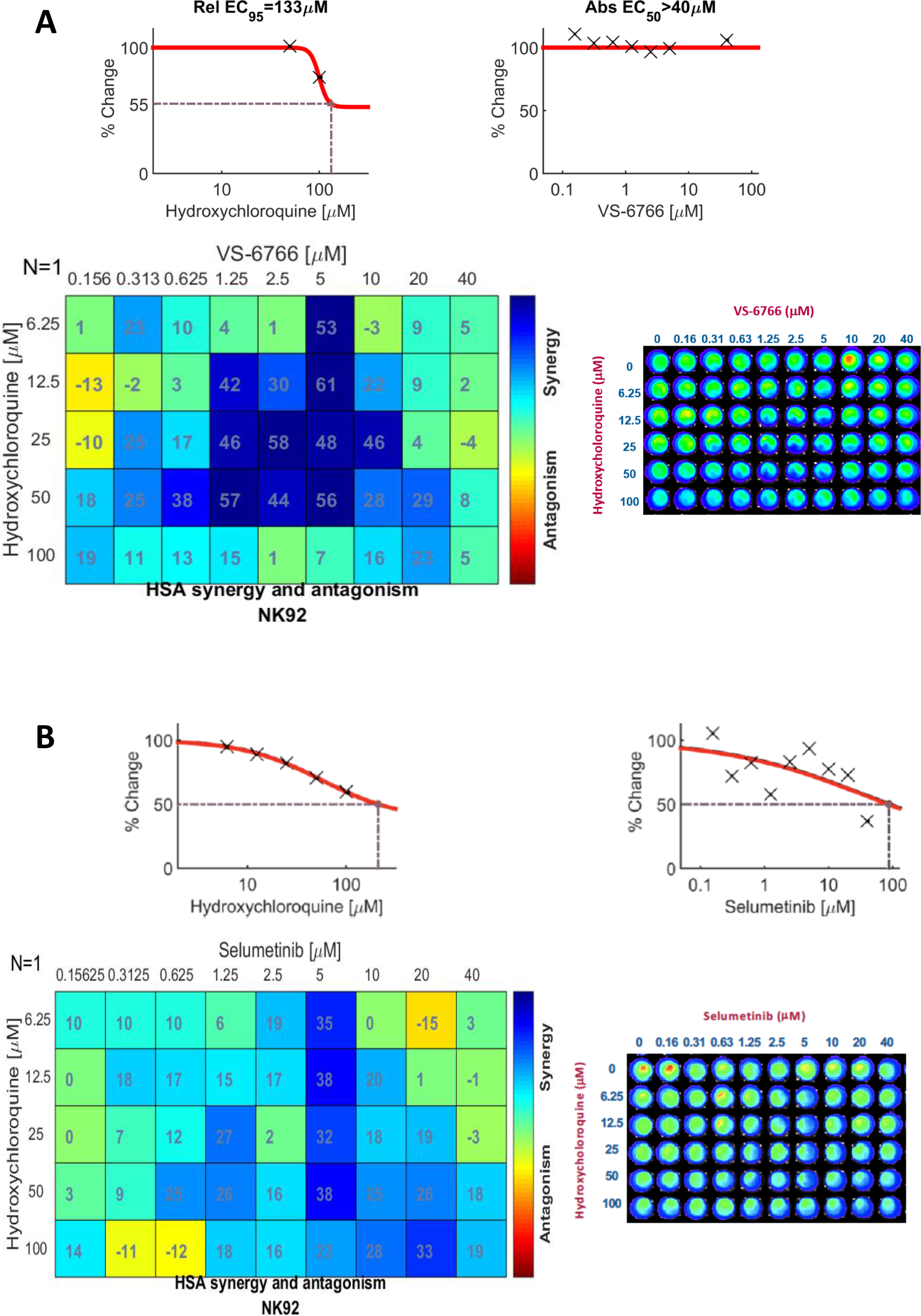
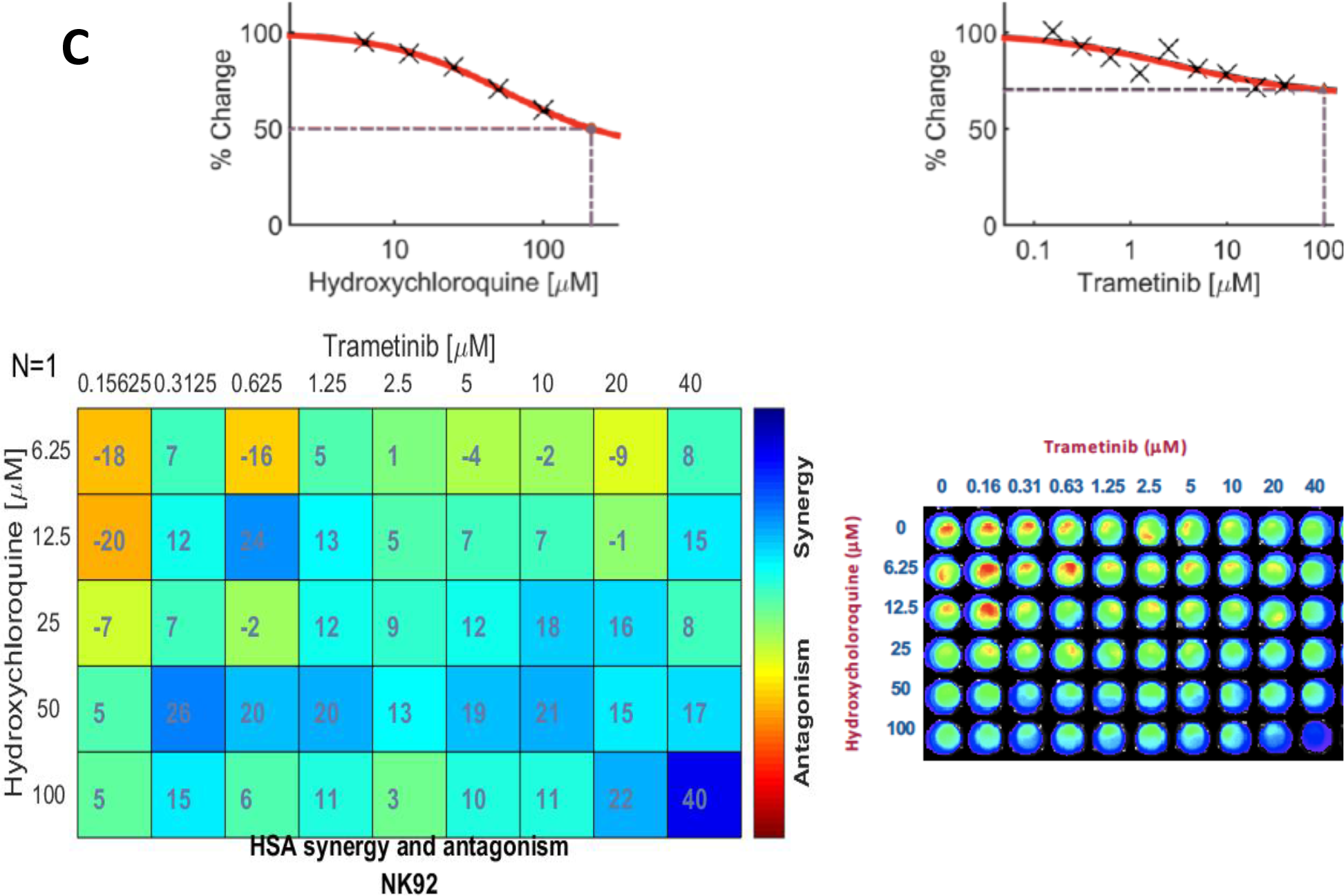
Cell viability assay of MRC-5 (A), H460 (B), H1299 (C) and BEAS-2B (D) treated with Chloroquine, Hydroxychloroquine and MEK inhibitors (VS-6766, Selumetinib, Trametinib and U0126). MRC-5 normal human lung fibroblast cells, H460 and H1299 human NSCLC cells, and BEAS-2B normal human bronchial airway epithelial cells were treated as indicated for 72 hours and CellTiter-Glo was added to acquire cell viability images with the IVIS system. GraphPad Prism 6 was used to plot the dose-response curve and calculate the IC50 values.

**Figure S9.**
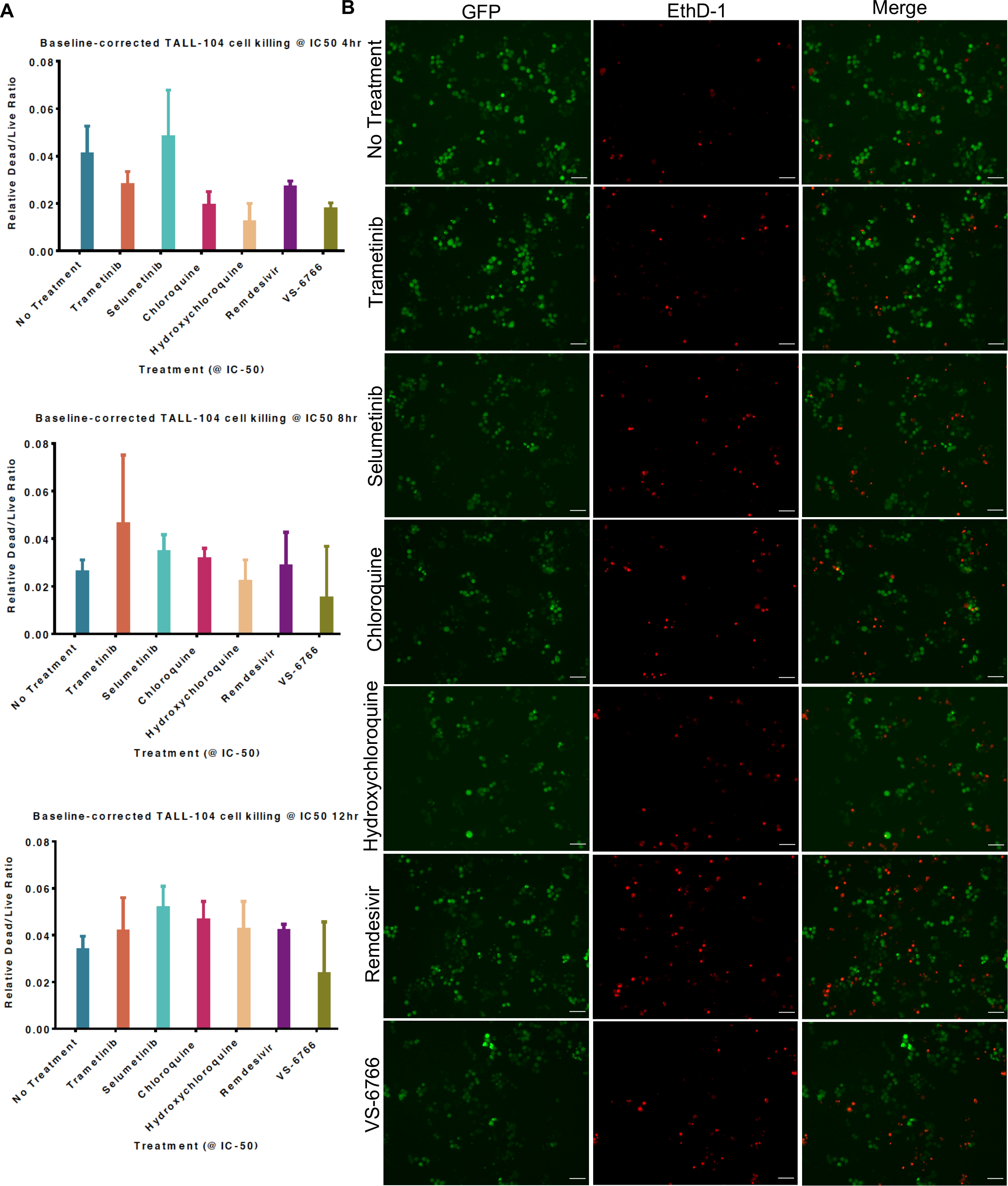
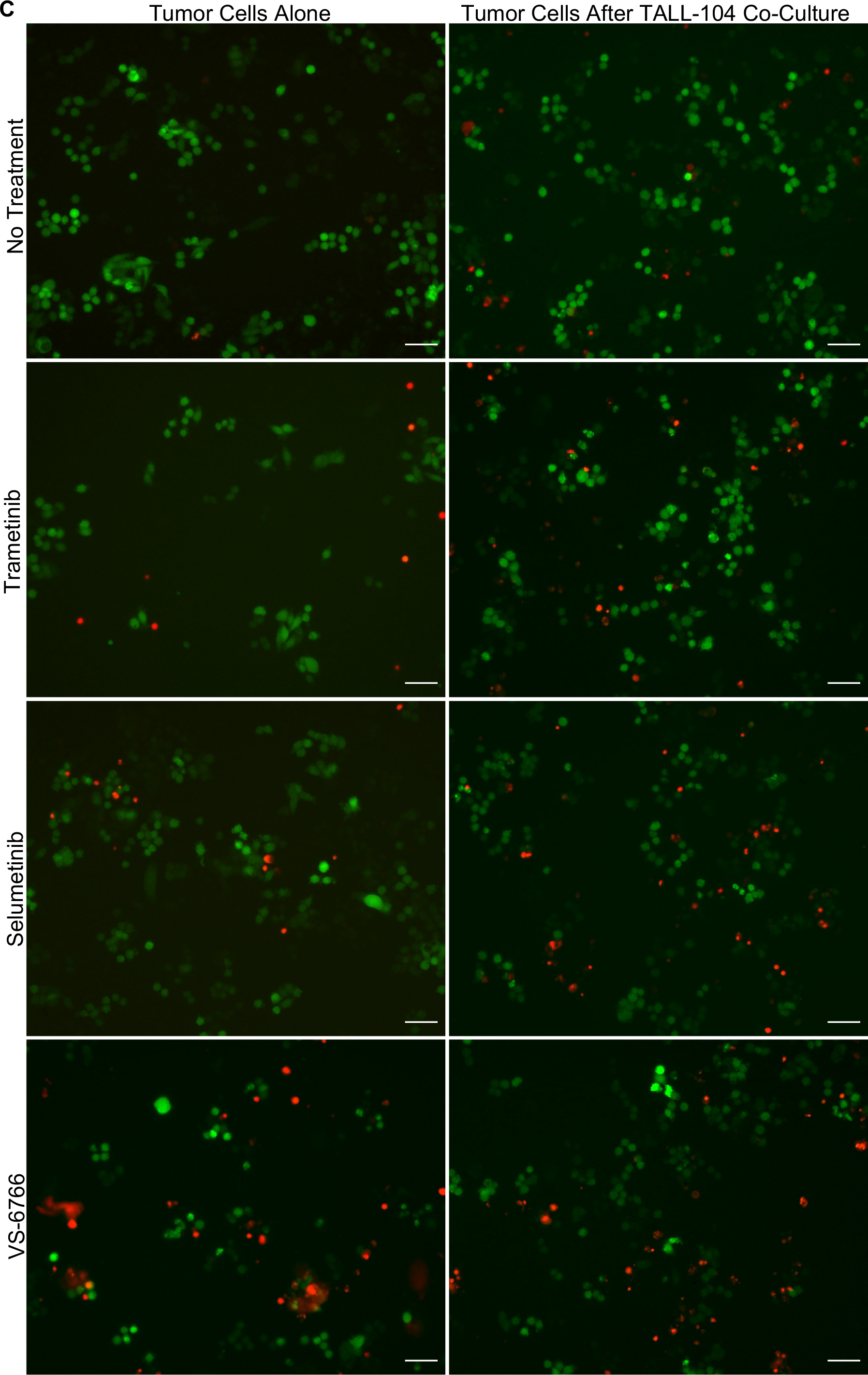
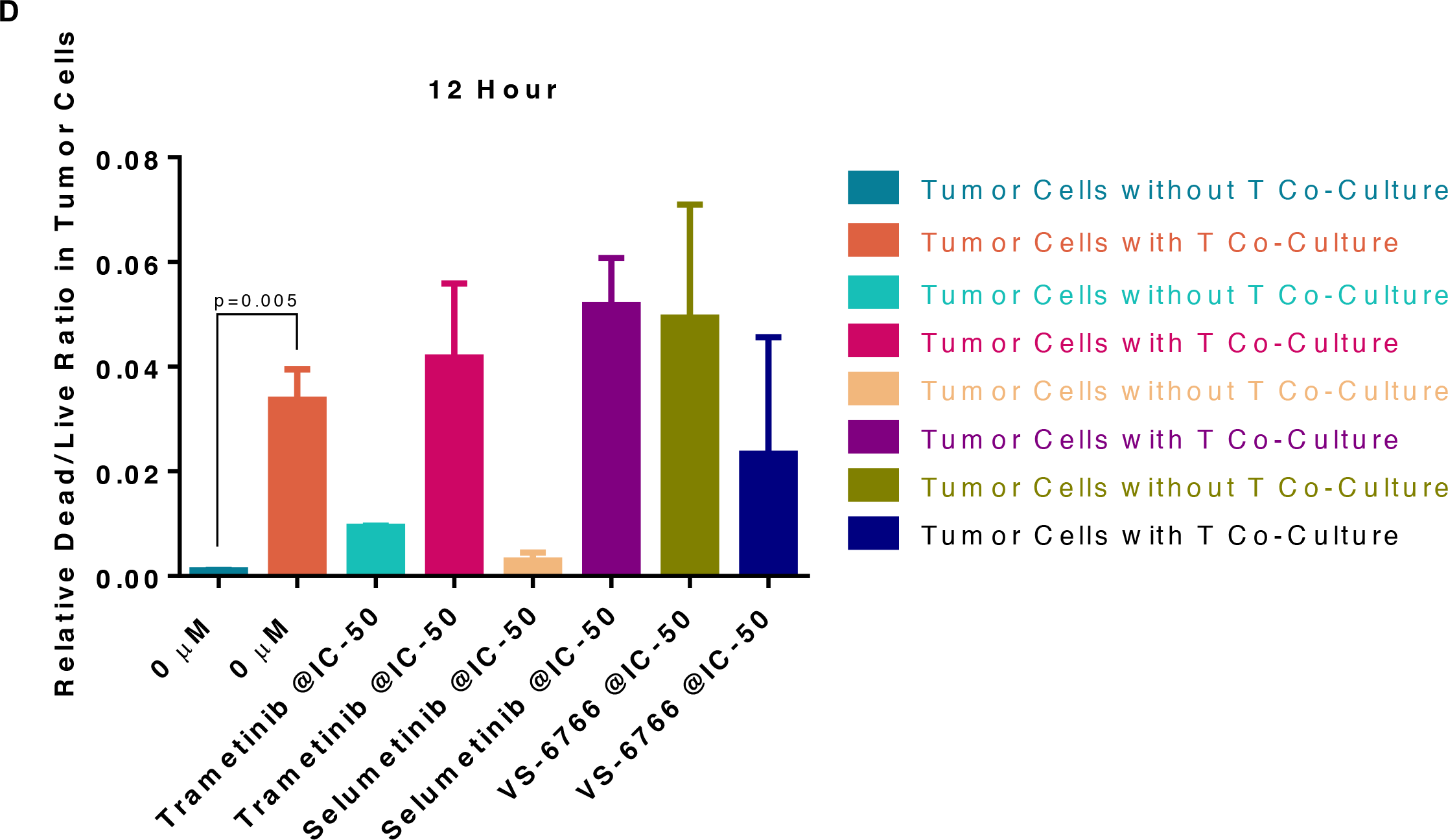
Combinational effect of Hydroxychloroquine and MEK inhibitors VS-6766 (A), Selumetinib (B) and Trametinib (C) on cell viability in NK-92 cells. NK-92 natural killer cells were treated as indicated for 72 hours and CellTiter-Glo was added to acquire cell viability images with the Xenogen IVIS system. Combenefit was used to plot single agent dose response and synergy distribution matrix.

**Figure S10.**
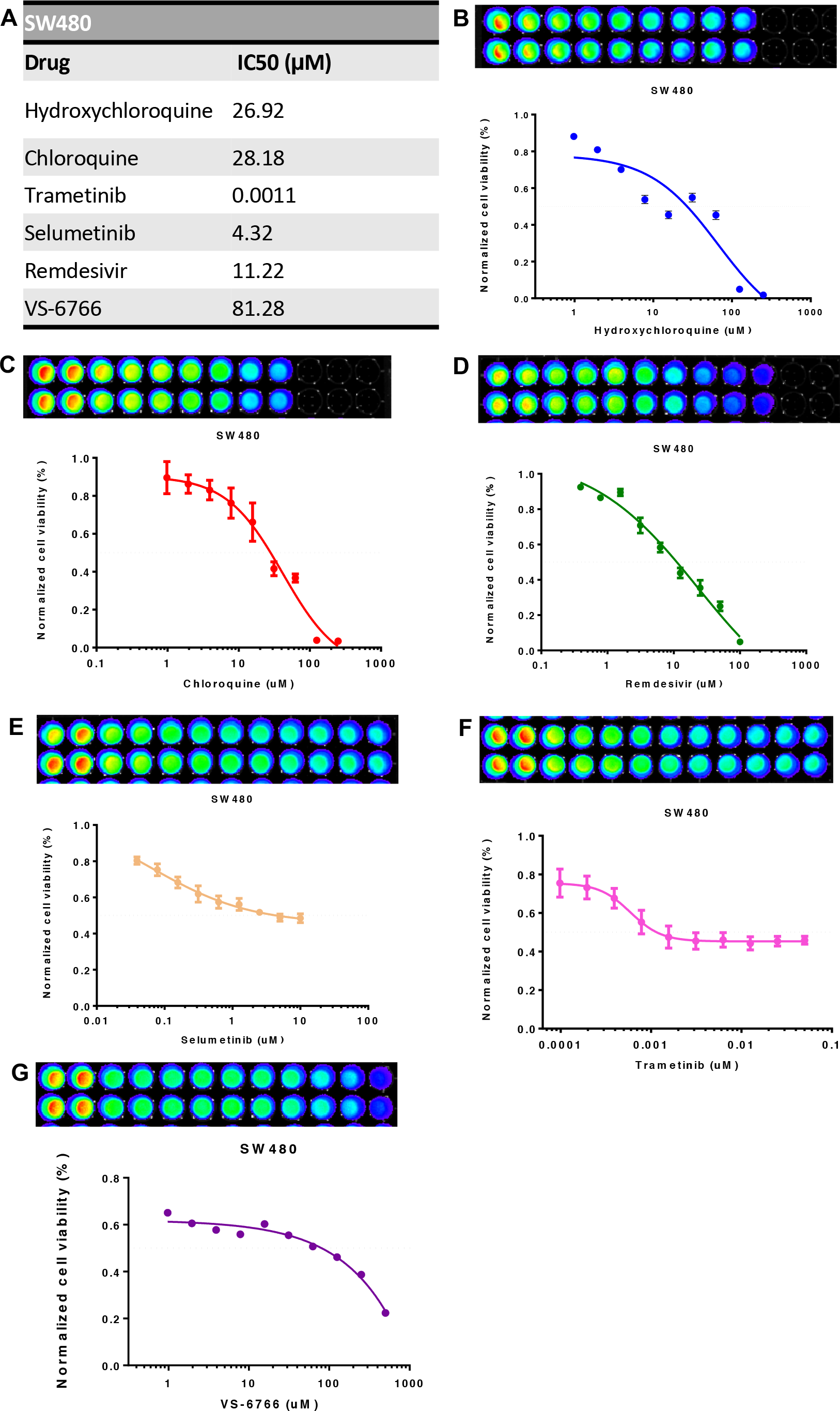
IC-50 values for SW480 cell line (A) as determined by cell viability assays of SW480 treated with Hydroxychloroquine (B), Chloroquine (C), Remdesivir (D), and MEK inhibitors Selumetinib, Trametinib, and VS-6766 (E-G). SW480 tumor cells were treated as indicated for 72 hours and CellTiter-Glo was added to acquire cell viability images with the IVIS system. GraphPad Prism 6 was used to plot the dose-response curve using a non-linear regression curve fit and to calculate the IC50 values.

**Figure S11.**
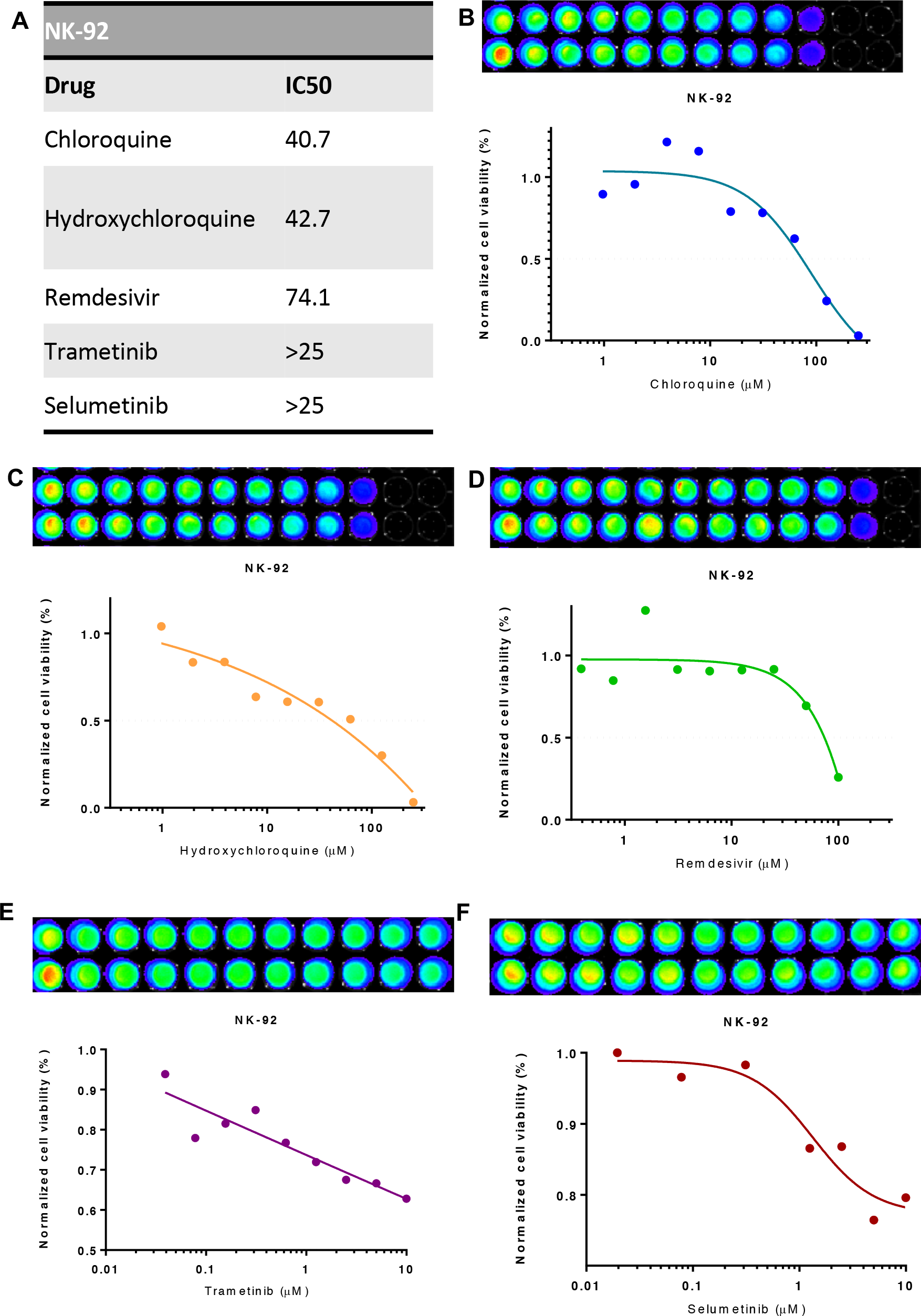
IC-50 values for NK-92 cell line (A) as determined by cell viability assays of NK-92 cells treated with Chloroquine (B), Hydroxychloroquine (C), Remdesivir (D), and MEK inhibitors Trametinib (E) and Selumetinib (F). NK-92 natural killer cells were treated as indicated for 72 hours and CellTiter-Glo was added to acquire cell viability images with the IVIS system. GraphPad Prism 6 was used to plot the dose-response curve using a non-linear regression curve fit and to calculate the IC50 values.

**Figure S12.**
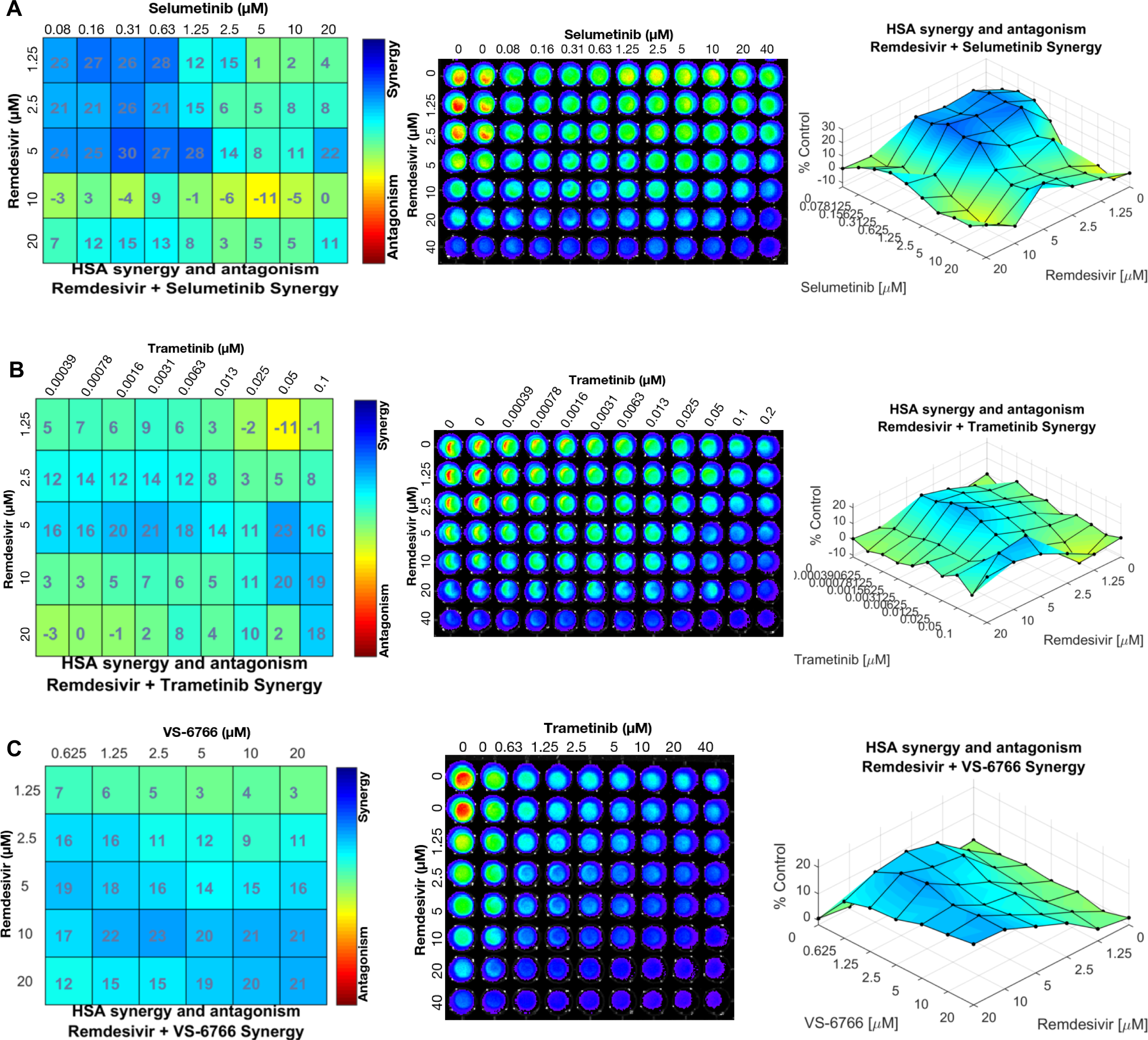
Combinational effect of Remdesivir and MEK inhibitors Selumetinib (A), Trametinib (B) and VS-6766 (C) on cell viability in SW480 tumor cells. SW480 tumor cells were treated as indicated for 72 hours and CellTiter-Glo was added to acquire cell viability images with the Xenogen IVIS system. Combenefit was used to plot synergy distribution matrices and combination dose response surface models.

## References

1. Guan, W.-j., et al., Clinical Characteristics of Coronavirus Disease 2019 in China. New England Journal of Medicine, 2020. 382(18): p. 1708–1720.

2. Goyal, P., et al., Clinical Characteristics of Covid-19 in New York City. New England Journal of Medicine, 2020. 382(24): p. 2372–2374.

3. McMichael, T.M., et al., Epidemiology of Covid-19 in a Long-Term Care Facility in King County, Washington. New England Journal of Medicine, 2020. 382(21): p. 2005–2011.

4. Guan, W.-j., et al., Comorbidity and its impact on 1590 patients with COVID-19 in China: a nationwide analysis. European Respiratory Journal, 2020. 55(5): p. 2000547.

5. Stawicki, S., et al., The 2019–2020 novel coronavirus (severe acute respiratory syndrome coronavirus 2) pandemic: A joint american college of academic international medicine-world academic council of emergency medicine multidisciplinary COVID-19 working group consensus paper. Journal of Global Infectious Diseases, 2020. 12(2): p. 47–93.

6. Kuderer, N.M., et al., Clinical impact of COVID-19 on patients with cancer (CCC19): a cohort study. The Lancet, 2020. 395(10241): p. 1907–1918.

7. Lee, L.Y.W., et al., COVID-19 mortality in patients with cancer on chemotherapy or other anticancer treatments: a prospective cohort study. The Lancet, 2020. 395(10241): p. 1919–1926.

8. Yang, K., et al., Clinical characteristics, outcomes, and risk factors for mortality in patients with cancer and COVID-19 in Hubei, China: a multicentre, retrospective, cohort study. The Lancet Oncology, 2020. 21(7): p. 904–913.

9. Bhatraju, P.K., et al., Covid-19 in Critically Ill Patients in the Seattle Region — Case Series. New England Journal of Medicine, 2020. 382(21): p. 2012–2022.

10. Mallapaty, S., Mini organs reveal how the coronavirus ravages the body. Nature, 2020. 583(7814): p. 15–16.

11. Zaim, S., et al., COVID-19 and Multiorgan Response. Current Problems in Cardiology, 2020. 45(8): p. 100618.

12. Ellul, M.A., et al., Neurological associations of COVID-19. The Lancet Neurology, 2020.

13. Varatharaj, A., et al., Neurological and neuropsychiatric complications of COVID-19 in 153 patients: a UK-wide surveillance study. The Lancet Psychiatry, 2020.

14. Ronco, C., T. Reis, and F. Husain-Syed, Management of acute kidney injury in patients with COVID-19. The Lancet Respiratory Medicine, 2020. 8(7): p. 738–742.

15. Becker, R.C., COVID-19 update: Covid-19-associated coagulopathy. J Thromb Thrombolysis, 2020. 50(1): p. 54–67.

16. Hoffmann, M., et al., SARS-CoV-2 Cell Entry Depends on ACE2 and TMPRSS2 and Is Blocked by a Clinically Proven Protease Inhibitor. Cell, 2020. 181(2): p. 271–280 e8.

17. Guaraldi, G., et al., Tocilizumab in patients with severe COVID-19: a retrospective cohort study. The Lancet Rheumatology, 2020.

18. Horby, P., et al., Effect of Dexamethasone in Hospitalized Patients with COVID-19: Preliminary Report. medRxiv, 2020: p. 2020.06.22.20137273.

19. Beigel, J.H., et al., Remdesivir for the Treatment of Covid-19 — Preliminary Report. New England Journal of Medicine, 2020.

20. Casadevall, A., M.J. Joyner, and L.-A. Pirofski, A Randomized Trial of Convalescent Plasma for COVID-19—Potentially Hopeful Signals. JAMA, 2020.

21. Konig, M.F., et al., Targeting the catecholamine-cytokine axis to prevent SARS-CoV-2 cytokine storm syndrome. medRxiv, 2020: p. 2020.04.02.20051565.

22. Riva, L., et al., Discovery of SARS-CoV-2 antiviral drugs through large-scale compound repurposing. Nature, 2020.

23. Bojkova, D., et al., Proteomics of SARS-CoV-2-infected host cells reveals therapy targets. Nature, 2020. 583(7816): p. 469–472.

24. Tay, M.Z., et al., The trinity of COVID-19: immunity, inflammation and intervention. Nature Reviews Immunology, 2020. 20(6): p. 363–374.

25. Korber, B., et al., Tracking Changes in SARS-CoV-2 Spike: Evidence that D614G Increases Infectivity of the COVID-19 Virus. Cell, 2020.

26. Leyfman, Y., et al., Potential Immunotherapeutic Targets For Hypoxia Due to COVI-FLU. Shock, 2020.

27. Hammer, Q., T. Rückert, and C. Romagnani, Natural killer cell specificity for viral infections. Nature Immunology, 2018. 19(8): p. 800–808.

28. Wu, G.S., et al., KILLER/DR5 is a DNA damage–inducible p53–regulated death receptor gene. Nature Genetics, 1997. 17(2): p. 141–143.

29. Carneiro, B.A. and W.S. El-Deiry, Targeting apoptosis in cancer therapy. Nat Rev Clin Oncol, 2020. 17(7): p. 395–417.

30. Wagner, J., et al., Dose intensification of TRAIL-inducing ONC201 inhibits metastasis and promotes intratumoral NK cell recruitment. J Clin Invest, 2018. 128(6): p. 2325–2338.

31. Sahin, I., et al., AMG-232 sensitizes high MDM2-expressing tumor cells to T-cell-mediated killing. Cell Death Discov, 2020. 6: p. 57.

32. Ricci, M.S., et al., Reduction of TRAIL-induced Mcl-1 and cIAP2 by c-Myc or sorafenib sensitizes resistant human cancer cells to TRAIL-induced death. Cancer Cell, 2007. 12(1): p. 66–80.

33. Finnberg, N., A.J. Klein-Szanto, and W.S. El-Deiry, TRAIL-R deficiency in mice promotes susceptibility to chronic inflammation and tumorigenesis. J Clin Invest, 2008. 118(1): p. 111–23.

34. Ralff, M.D. and W.S. El-Deiry, TRAIL pathway targeting therapeutics. Expert Review of Precision Medicine and Drug Development, 2018. 3(3): p. 197–204.

35. Waggoner, S.N., et al., Roles of natural killer cells in antiviral immunity. Curr Opin Virol, 2016. 16: p. 15–23.

36. Masselli, E., et al., NK cells: A double edge sword against SARS-CoV-2. Advances in Biological Regulation, 2020. 77: p. 100737–100737.

37. Smyth, M.J., et al., Tumor Necrosis Factor–Related Apoptosis-Inducing Ligand (Trail) Contributes to Interferon γ–Dependent Natural Killer Cell Protection from Tumor Metastasis. Journal of Experimental Medicine, 2001. 193(6): p. 661–670.

38. Sato, K., et al., Antiviral response by natural killer cells through TRAIL gene induction by IFN–α/β European Journal of Immunology, 2001. 31(11): p. 3138–3146.

39. Chen, I.-Y., et al., Upregulation of the Chemokine (C-C Motif) Ligand 2 via a Severe Acute Respiratory Syndrome Coronavirus Spike-ACE2 Signaling Pathway. Journal of Virology, 2010. 84(15): p. 7703–7712.

40. Gallagher, P.E., C.M. Ferrario, and E.A. Tallant, MAP kinase/phosphatase pathway mediates the regulation of ACE2 by angiotensin peptides. Am J Physiol Cell Physiol, 2008. 295(5): p. C1169–74.

41. Goldman, J.D., et al., Remdesivir for 5 or 10 Days in Patients with Severe Covid-19. New England Journal of Medicine, 2020.

42. Fukushi, S., et al., Vesicular stomatitis virus pseudotyped with severe acute respiratory syndrome coronavirus spike protein. J Gen Virol, 2005. 86(Pt 8): p. 2269–2274.

43. Kobinger, G.P., et al., Human immunodeficiency viral vector pseudotyped with the spike envelope of severe acute respiratory syndrome coronavirus transduces human airway epithelial cells and dendritic cells. Hum Gene Ther, 2007. 18(5): p. 413–22.

44. Fukuma, A., et al., Inability of rat DPP4 to allow MERS-CoV infection revealed by using a VSV pseudotype bearing truncated MERS-CoV spike protein. Arch Virol, 2015. 160(9): p. 2293–300.

45. Bouhaddou, M., et al., The Global Phosphorylation Landscape of SARS-CoV-2 Infection. Cell, 2020.

46. Cao, W. and T. Li, COVID-19: towards understanding of pathogenesis. Cell Res, 2020. 30(5): p. 367–369.

47. Wiersinga, W.J., et al., Pathophysiology, Transmission, Diagnosis, and Treatment of Coronavirus Disease 2019 (COVID-19): A Review. JAMA, 2020.

48. Vijayvargiya, P., et al., Treatment Considerations for COVID-19: A Critical Review of the Evidence (or Lack Thereof). Mayo Clin Proc, 2020. 95(7): p. 1454–1466.

49. Dieterle, M.E., et al., A replication-competent vesicular stomatitis virus for studies of SARS-CoV-2 spike-mediated cell entry and its inhibition. bioRxiv, 2020.

50. Case, J.B., et al., Neutralizing antibody and soluble ACE2 inhibition of a replication-competent VSV-SARS-CoV-2 and a clinical isolate of SARS-CoV-2. Cell Host & Microbe, 2020.

51. Smith, J.A., et al., Suppression of mitochondrial biogenesis through toll-like receptor 4-dependent mitogen-activated protein kinase kinase/extracellular signal-regulated kinase signaling in endotoxin-induced acute kidney injury. J Pharmacol Exp Ther, 2015. 352(2): p. 346–57.

52. Smith, J.A., P.R. Mayeux, and R.G. Schnellmann, Delayed Mitogen-Activated Protein Kinase/Extracellular Signal-Regulated Kinase Inhibition by Trametinib Attenuates Systemic Inflammatory Responses and Multiple Organ Injury in Murine Sepsis. Crit Care Med, 2016. 44(8): p. e711–20.

53. Kurian, N., et al., Dual Role For A MEK Inhibitor As A Modulator Of Inflammation And Host Defense Mechanisms With Potential Therapeutic Application In COPD. Int J Chron Obstruct Pulmon Dis, 2019. 14: p. 2611–2624.

54. Au, E.D., et al., The MEK-Inhibitor Selumetinib Attenuates Tumor Growth and Reduces IL-6 Expression but Does Not Protect against Muscle Wasting in Lewis Lung Cancer Cachexia. Front Physiol, 2016. 7: p. 682.

55. Baumann, D., et al., Proimmunogenic impact of MEK inhibition synergizes with agonist anti-CD40 immunostimulatory antibodies in tumor therapy. Nat Commun, 2020. 11(1): p. 2176.

